# Minicollagen expression dynamics reveal a transcriptional program for cnidogenesis in the sea anemone *Nematostella vectensis*

**DOI:** 10.64898/2026.06.23.733813

**Authors:** Anna M. L. Klompen, Jenny Duong, Mary C. McKinney, Jason A. Morrison, Jose E. Javier, Shiyuan Chen, Sean McKinney, Kate E. Hall, Kaitlyn Petentler, Lacey Ellington, Matthew C. Gibson

**Affiliations:** Stowers Institute for Medical Research, Kansas City, MO, USA

**Keywords:** nematocyte, spirocyte, nematogenesis, spirogenesis, HCR RNA-FISH, RNA-FACS-seq, differential gene expression

## Abstract

Cnidae are explosive harpoon-like organelles localized within stinging cells, or cnidocytes, of the phylum Cnidaria (jellyfish, hydroids, sea anemones, and corals). These unique Golgi-derived structures define the phylum and are prominent examples of evolutionary novelty at the subcellular level. While recent studies have focused on the developmental specification of cnidocytes, less is known about the genetic basis for structural variation and differential toxin expression between distinct cnidae subtypes. Here we determine the transcriptional profile of developing nematocytes and spirocytes, the two major cnidae subtypes in the sea anemone *Nematostella vectensis*. We first interrogate the gene expression patterns and cell type-specificity of three known and three uncharacterized minicollagen genes, which encode a family of cnidae-specific structural proteins. We report that four minicollagen genes are broadly expressed in ectodermal cnidocytes while two others are restricted to tentacular cnidocytes. Leveraging whole adult scRNA-seq analysis as well as nematocyte- and spirocyte-specific transgenic reporter lines, we then identify the tentacle-restricted cnidocytes as spirocytes distinguished by expression of the minicollagen *NvNcol5*. To deepen our understanding of the gene expression differences between nematocytes and spirocytes, we next developed a customized RNA-FACS-seq pipeline. This approach identified a suite of differentially expressed genes, illuminating spatial and temporal gene expression dynamics in both cell types. Altogether, our experiments provide fundamental and novel insights into the specialization of cnidarian stinging cells while establishing a rich set of molecular resources for further investigation.

## INTRODUCTION

Exploring the genetic basis for the specification and morphogenesis of novel cell types can inform our understanding of macroevolutionary shifts and adaptation^1–6^. Cnidocytes or “stinging cells,” are the defining feature of the phylum Cnidaria (jellyfish, sea anemones, and corals), and are critical for predation, defense, and for some species intraspecific competition, settlement, and locomotion^7,8^. These cells produce explosive Golgi-secreted organelles known as cnidae or cnidocysts^8^. Cnidae are well established as cellular novelties^9^, and have been described as among the most complex subcellular structures in the animal kingdom^7^. The intricate morphologies of cnidae are readily visible by light microscopy, and at least 25-30 different morphological types have been described across cnidarians with variable levels of taxonomic specificity^8,10,11^. Broadly, there are three major categories of cnidae: nematocysts, spirocysts, and ptychocysts^10–13^. Nematocysts are found throughout the phylum and there are typically multiple distinct morphological types within an individual^8,9,11,14^. Nematocysts often contain toxin mixtures (venoms) used for prey capture or deterring predators^15^, with different nematocyst types deploying distinct venom components^16^. Unlike the ubiquitous nematocytes, spirocysts and ptychocysts are restricted to specific clades within the class Anthozoa (corals and sea anemones) and are not thought to deploy venom^12,13^. Spirocysts are instead used to entangle prey and have a thinner capsule wall with a tightly coiled tubule of uniform diameter^12^. Even more specialized, ptychocysts are used to build the characteristic tube-dwelling structure of species in the order Ceriantharia^13^. The majority of cnidae diversity is thus observed in nematocysts, while spirocysts and ptychocysts are more morphologically constrained^8,11^. However, the molecular basis for this morphological variation remains poorly characterized, despite the rising number of single-cell based datasets from a variety of cnidarian species^17–31^. Accordingly, our ability to discern major cnidocyte types and subtypes and to interrogate their development remains challenging^17,18,22–25,28,30,31,32^.

The starlet sea anemone *Nematostella vectensis* (Stephenson, 1935)^33^, has been the focus of multiple molecular studies on cnidocyte biology^9,31,34–43^. *Nematostella* polyps possess spirocysts and three morphologically variant types of nematocysts^43,44^. The number and localization of these cnidocyte types vary from early development to mature adults (**Figure 1**). For example, nematocytes appear during early life stages and remain abundant across the ectoderm throughout development, though several specific subtypes emerge within tissues that develop at later life stages. Spirocytes are typically only observed in the tentacles, which are most prominent in juvenile and adult polyps (**Figure 1B-C**). Due to the scarcity of this cell type during early development, only a few spirocyte-specific molecular markers have been established, including *Calumenin F* (*NvCaluF*; NV2.16293^38^; previously called *SpCARP1-like*^44^) and *FoxL2* (NV2.14459)^17,31^, though *FoxL2* also labels a population of neurons^17,46^,. While there are several scRNA-seq datasets that include putative spirocyte clusters^17–19^, few markers have been characterized *in situ*.

**Figure 1:**
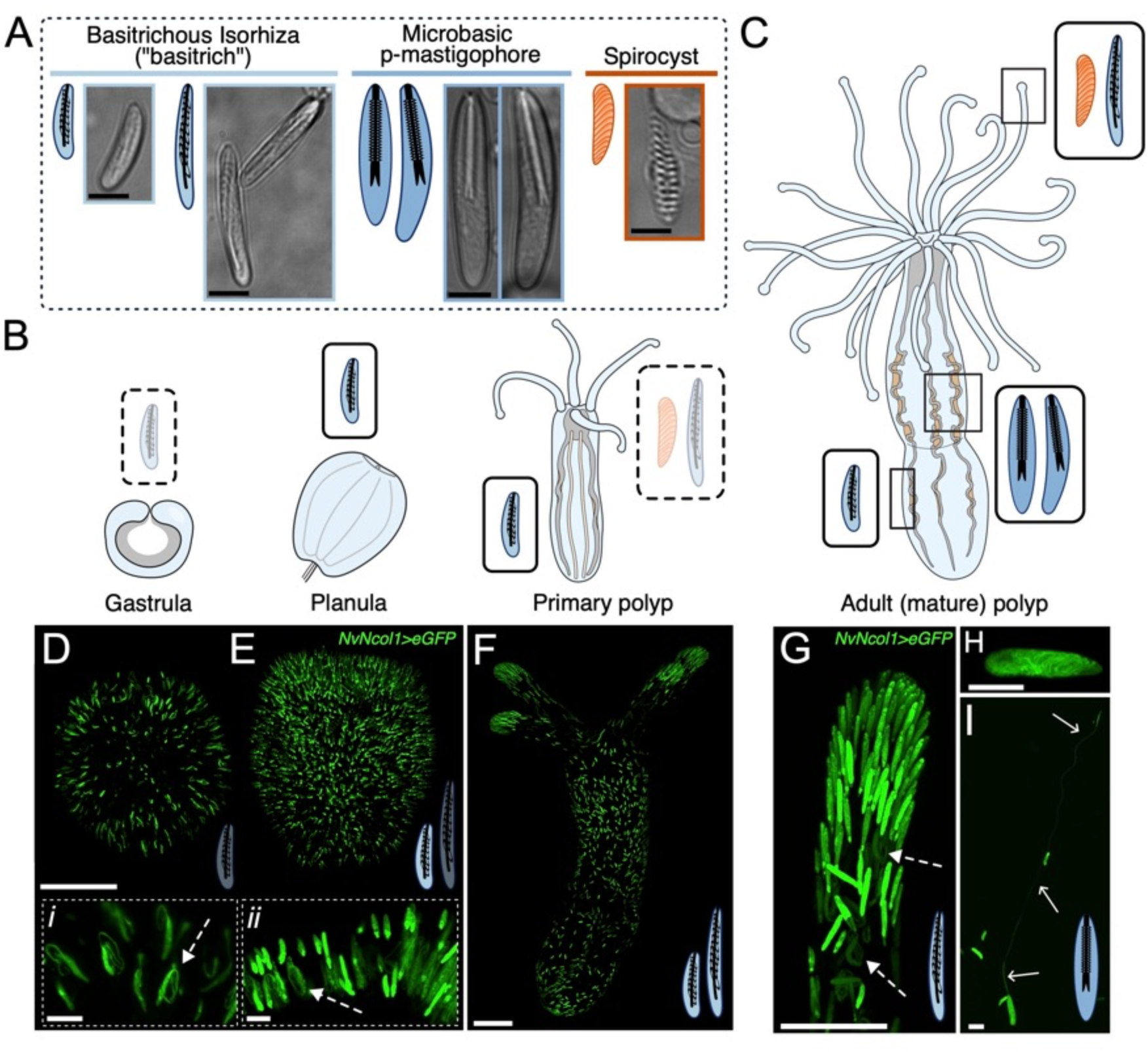
Cnidae diversity and development across major life stages of *Nematostella*. **A)** Four major cnidae morphologies observed in *Nematostella*, with nematocyst subtypes highlighted in blue and spirocysts highlighted in orange. **B)** Schematic of late gastrula, planula, primary polyp, and **C)** adult stage of *Nematostella*. Drawings are modified illustrations from Mark Miller. Boxes contain relevant cnidae morphologies at each stage, where dotted boxes indicate cnidocyte subtypes currently developing. Boxes along the adult tissue indicate where each cnidocyte type is at the highest proportion, in accordance with Zenkert *et al.* ^43^; spirocytes and large isorhize in tentacles, small isorhizae along ectoderm of the body column, and mastigophores lining the internal mesenteries. **D)** Early planula (2.5dpf), **E)** late planula (5dpf), and **F)** juvenile primary polyp (10dpf) from *NvNcol1>eGFP^cnidae^* line. All animals were raised at 17C. Schematics of cnidae indicate types present at each stage, with translucent cnidae indicating that type is under active development. eGFP from *NvNcol1>eGFP^cnidae^* is actively trafficked into the developing capsule and tubule during nematogenesis, such that the organelle is visible through mid-development and recent maturation. Insets in **D)** and **E)** indicate higher magnification from **inset i)** early planula and **inset ii)** late planula, respectively. White dotted arrows indicate developing cnidae, with tubule forming on outside of capsule. **G-I)** Live images from *NvNcol1>eGFP^cnidae^* line from adult tissues. **G)** Tentacle tip with majority of eGFP+structures are corresponding with large isorhizae type nematocysts, with those under development indicated by white dotted arrows. Spirocysts are present in this tissue but do not express eGFP. **H)** A mature mastigophore capsule, with inverted tubule clearly visible. **I)** Discharged mastigophore, with extroverted shaft and tubule (white arrows) maintaining eGFP signal. Scale bars: **A)** 5 μm, **D-F)** 100 μm, **insets** 10 μm, **G)** 50 μm, **H-I)** 10 μm.

In *Nematostella*, a family of structural proteins known as minicollagens (NvNcols, also called NvMcols) are known to exhibit cnidae type-specific expression^43^. Minicollagens function as structural scaffolding for developing cnidae capsules, in some cases, the spined tubules^47^. Minicollagen genes have thus been employed as markers for cnidae biogenesis (cnidogenesis) and further leveraged in the production of cnidocyte-specific reporter lines^14,48,49^, including *NvNcol3* in *Nematostella*^34^. Five minicollagens have been previously identified in *Nematostella*, with three characterized by *in situ* hybridization and immunohistochemistry^43^. *NvNcol1* (NV2.12621) is only observed in developing nematocytes, while *NvNcol3* (NV2.10686) and *NvNcol4* (NV2.11801) are described as universal cnidocyte markers that are present in both nematocytes and spirocytes. However, scRNA-seq datasets have provided more expansive, and somewhat conflicting, evidence for *Nematostella* minicollagen expression across different cnidocyte types. Therefore, a systematic experimental validation of minicollagen gene expression dynamics would provide an important molecular entry point for the downstream analysis of spirocyte biology and cnidogenesis more broadly.

In the present work we clarify and then leverage the dynamic expression patterns of minicollagens to uncover the molecular programs that distinguish developing nematocytes and spirocytes in *Nematostella*. By integrating multiplexed *in situ* experiments, adult whole-animal scRNA-seq and generation of our own transgenic lines, we define the cell type-specific gene expression patterns of six *Nematostella* minicollagens across cnidogenesis. These analyses demonstrate that two minicollagens, *NvNcol5* and another defined herein as *NvNcol0* (previously *Ncol*^17–19,42,50^), are specifically expressed in developing spirocytes. We further develop a customized RNA-FACS-seq protocol^51–54^ to isolate fixed cell populations expressing each minicollagen for bulk RNA-sequencing (RNA-seq). Using this approach, we determine the global gene expression dynamics that distinguish spirocytes and nematocytes. Combined, these datasets provide a detailed window into the molecular profiles of two major cnidocyte lineages and establish foundational resources for the functional exploration of these two novel cell types.

## RESULTS

### Generation of a nematocyte-specific reporter line in Nematostella

To visualize nematocytes during *Nematostella* development, we first established a novel fluorescent reporter line using the putative promoter for *Nematostella minicollagen 1* (*NvNcol1>eGFP*^cnidae^; **Supplemental Data S1**). As expected, eGFP was observed in developing and recently matured nematocytes, including small isorhizae across all life stages and large isorhizae and microbasic p-mastigophores at the polyp stage (**Figure 1A, D-I**). Within 24 hours post fertilization (hpf) at room temperature, eGFP+ nematocytes were observed throughout the ectoderm of developing planula larvae. At ∼3-4dpf we observed a high density of tentacle bud-associated large isorhizae developing at the oral pole (**Figure 1E**). eGFP+ cells were present throughout the ectoderm into adulthood, with a notably higher density in the tentacles (**Figure 1F,G).**

Inadvertently, the promoter region we used in *NvNcol1>eGFP*^cnidae^ contained a portion of the *NvNcol1* gene body (**Supplemental Data S1**). This serendipitously resulted in the subcellular localization of eGFP to both the capsule and intricate tubule structure of developing nematocysts. We were thus able to directly visualize the formation and maturation of nematocysts *in vivo*, particularly the initiation, formation and inversion of the tubule. We documented nematogenesis throughout the ectoderm of early (**Figure 1D, inset *i***) and late-stage planulae (**Figure 1E, inset *ii***), as well as in the tentacles of adult animals (**Figure 1G, arrow**). In high resolution images, eGFP fluorescence localized to the tubule in both mature (**Figure 1H**) and discharged nematocytes (**Figure 1I, arrows**). To ensure the inclusion of the gene body did not impact the overall expression pattern, we development a second reporter by isolating only the putative promoter (*NvNcol1>eGFP*) and validated that eGFP+ nematocytes were expressed in developing and mature nematocytes (**Supplemental Fig. S1).** Overall, analysis of these new transgenic lines showed that *NvNcol1* is a robust nematocyte-specific marker throughout the life cycle of *Nematostella*. Based on the localization of eGFP within developing cnidae in *NvNcol1>eGFP*^cnidae^, we further conclude that *NvNcol1* encodes both a structural component of the capsule and, most likely, the tubule as well.

### Nematostella minicollagens display cell type-specific gene expression dynamics

After validating *NvNcol1* expression using our reporter line, we next employed multiplexed hybridization chain reaction RNA-FISH (HCR) to determine the correlated gene expression profiles for each of the six *Nematostella* minicollagen genes. Specifically, we monitored the expression of *NvNcol0* (NV2.11820)*, NvNcol1*, *NvNcol3*, *NvNcol4*, *NvNcol5* (NV2.11802), and *NvNcol6* (NV2.11815) and in early planulae (2dpf), budding planulae (4dpf), and primary polyps (5dpf). Each minicollagen gene displayed robust signal and exhibited a distinct expression pattern (**Figure 2**; amplifier-only controls shown in **Supplemental Fig. S2**). As expected, both *NvNcol3*+ (**Figure 2B**) and *NvNcol1+* (**Figure 2E**) cells were observed throughout the ectoderm at all three stages, including a relatively high concentration within the tips of developing tentacles. *NvNcol4*+ cells were found throughout the ectoderm and typically overlapped with *NvNcol3+/NvNcol1+* cells. *NvNcol6+* cells were also broadly ectodermal and displayed consistent overlap with *NvNcol3*, *NvNcol1*, and *NvNcol4*. Interestingly, we observed a subset cells that expressed solely *NvNcol6*, often with a discernible ovular structure indicative of the later-stage capsule of a developing cnidocyte (**Figure 2A, *inset iii, iv***). This could indicate that *NvNcol6* is expressed at a later stage of cnidocyte development than cells expressing *NvNcol3*, *NvNcol1*, and *NvNcol4*. Overall, we found that *NvNcol4* and *NvNcol6* were consistently expressed with *NvNcol1*, suggesting they are specific to developing nematocytes, as well as some evidence of temporally ordered expression throughout nematogenesis (**Figure 2**).

**Figure 2.**
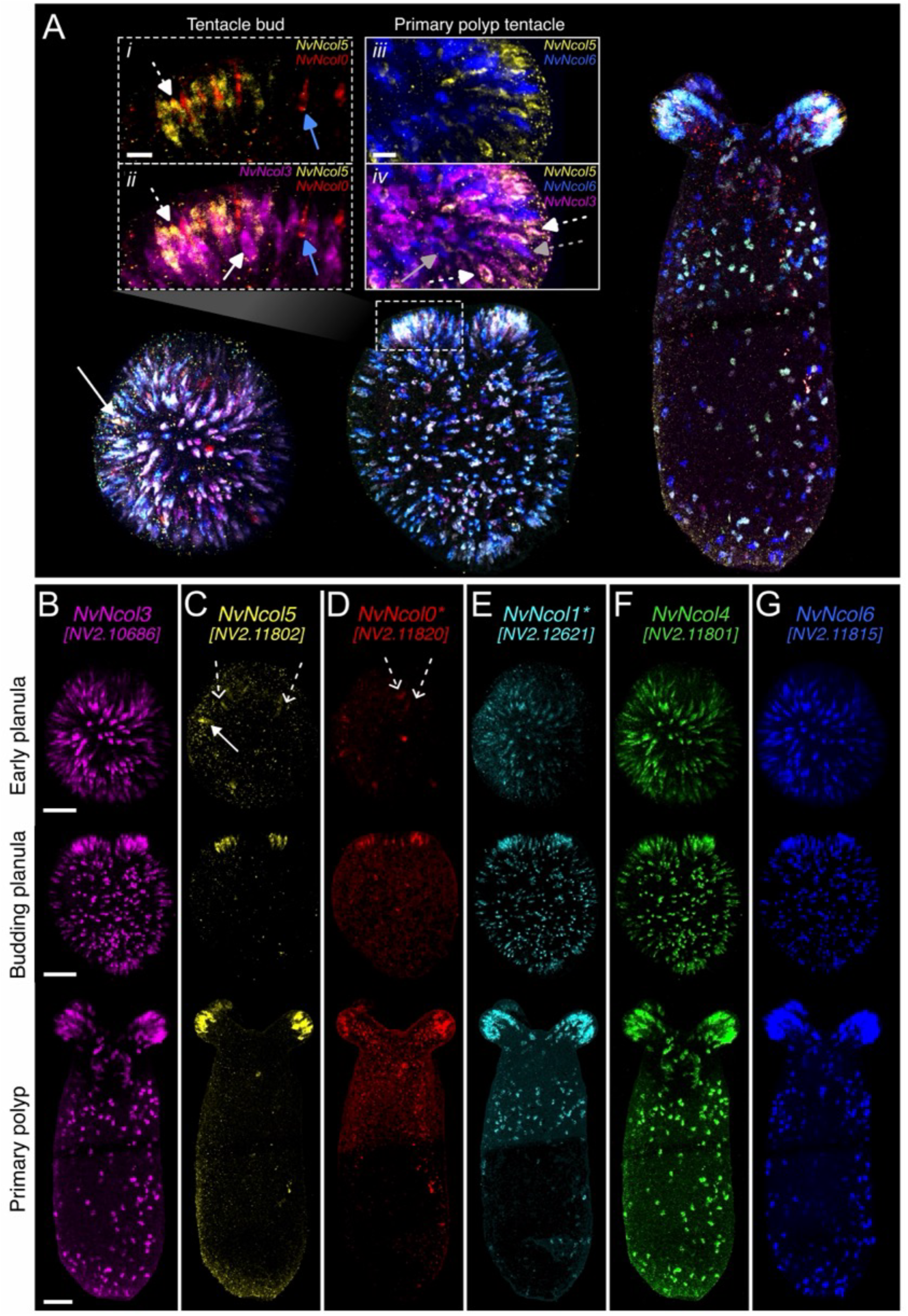
Expression of minicollagen gene family members during early developmental stages. **A)** Imaging from a 6-gene *minicollagen* multiplex HCR experiment for whole tissue from early planula (2dpf), budding planula (4dpf), and primary polyp (5dpf), raised at RT with oral end facing upwards. **Inset i)** Higher magnification images of *NvNcol5+* and *NvNcol0+* cells from the budding tentacle of the planula, where the white dotted arrow indicates overlap of the two genes while the blue solid arrow indicated a cell expressing only *NvNcol0*. **Inset ii)** Same image as **inset i**, but with the addition of *NvNcol3+* cells, a known marker for cnidogenesis. White dotted arrow and solid blue arrow indicate the same cells as previously, while the white solid arrow indicates a cell expressing only *NvNcol3*. **Inset iii)** High magnification image from tentacle tip of a primary polyp suggesting a lack of overlap between *NvNcol5+* cells and *NvNcol6+* cells. **Inset iv)** Same image as **inset iii**, but with addition of *NvNcol3+* cells, where the white solid arrows indicated overlap with *NvNcol5+* and gray solid arrows indicated overlap with *NvNcol6+*. The gray solid arrow indicates that some cells are *NvNcol6+* only. **B-G)** Gene-specific signal of each minicollagen from tissues represented in **A)**. White arrow in early planula of **A)** and **C)** point to the same putative cells, which appear to be *NvNcol5+* with relatively high signal. Dotted arrows in early planula for **C)** and **D)** show positive but weaker HCR signal. These positive cells suggest that cells specific to tentacle tissue are already developing before observing morphological changes associated with tentacle development. * indicate images that were split using lifetime unmixing. Scale bars: **A)** insets: 10 μm, **B-G)** 50 μm.

A further goal of our analysis was to evaluate the gene expression dynamics of *NvNcol5* and *NvNcol0*, which were identified as putative spirocyte-specific markers in previous scRNA-seq datasets^17–19^. Intriguingly, like spirocytes, cells expressing *NvNcol5* and *NvNcol0* were clearly restricted to the budding tentacles of late planula-stage animals as well as the tentacle tips of primary polyps (**Figures 2C and D**). High resolution co-expression analysis of *NvNcol5* and *NvNcol0* together with the cnidocyte marker *NvNcol3* revealed dynamic and overlapping patterns for all three genes. We found that *NvNcol5+* cells were more abundant than *NvNcol0+* cells, but these populations typically overlapped. A smaller subset of cells solely expressed *NvNcol0* (**Figure 2A, inset *i*)**. We also observed consistent co-expression of *NvNcol3* and *NvNcol5*, indicating these cells are indeed cnidocytes, while a subset of *NvNcol0+* cells lacked *NvNcol3* expression (**Figure 2A, inset *ii*)**. These cells consistently displayed an ovular capsule-like structure characteristic of more mature cnidocytes; we thus infer that *NvNcol0* could be expressed late in cnidogenesis at a stage when *NvNcol3* expression is reduced or absent. Collectively, these data are consistent with the interpretation that *NvNcol5* and *NvNcol0* reflect early and late spirocyte markers, respectively.

To directly distinguish between nematocytes and putative spirocytes *in vivo*, we next examined cells co-expressing *NvNcol3* (general cnidocyte marker), *NvNcol5* (putative spirocytes), and *NvNcol6* (nematocytes). In the tentacle tips of primary polyps, we found that *NvNcol3* overlapped with both *NvNcol5* and *NvNcol6.* However, we did not detect co-expression of *NvNcol5* and *NvNcol6* (**Figure 2A, inset *iii, iv*),** suggesting they mark two distinct cnidocyte populations. We conclude that *NvNcol5,* and subsequently *NvNcol0*, are expressed in a distinct tentacular cnidocyte type not labeled by nematocyte-specific minicollagens (e.g. *NvNcol1*, *NvNcol6*). More broadly, our multiplex HCR experiments confirm and extend the minicollagen gene expression profiles suggested by previous scRNA-seq datasets: *NvNcol3* is a general marker for all cnidocyte types while *NvNcol1*, *NvNcol4,* and *NvNcol6* broadly overlapped in developing nematocytes. In contrast, *NvNcol5* and *NvNcol0* were restricted to putative spirocytes in the tentacles.

### Minicollagen expression dynamics in adult cnidocyte cell lineages

To complement our analysis of *Nematostella* minicollagen gene expression during early development, we next employed scRNA-seq to generate a multi-individual, whole-adult cell atlas. We integrated datasets collected from both *wild-type* adults and FACS-enriched cells from a *NvNematogalectin>mScarlet-I* ^35^ reporter line to enrich for developing cnidocytes. Combined, we isolated 20,726 cells from six adult males (**Figure 3; Supplemental Figure S5**). From this dataset we identified the majority of previously characterized *Nematostella* cell types based on known markers and those established in Cole *et al.* ^19^ (**Supplemental Fig. S6, Supplemental Data S2**).

**Figure 3:**
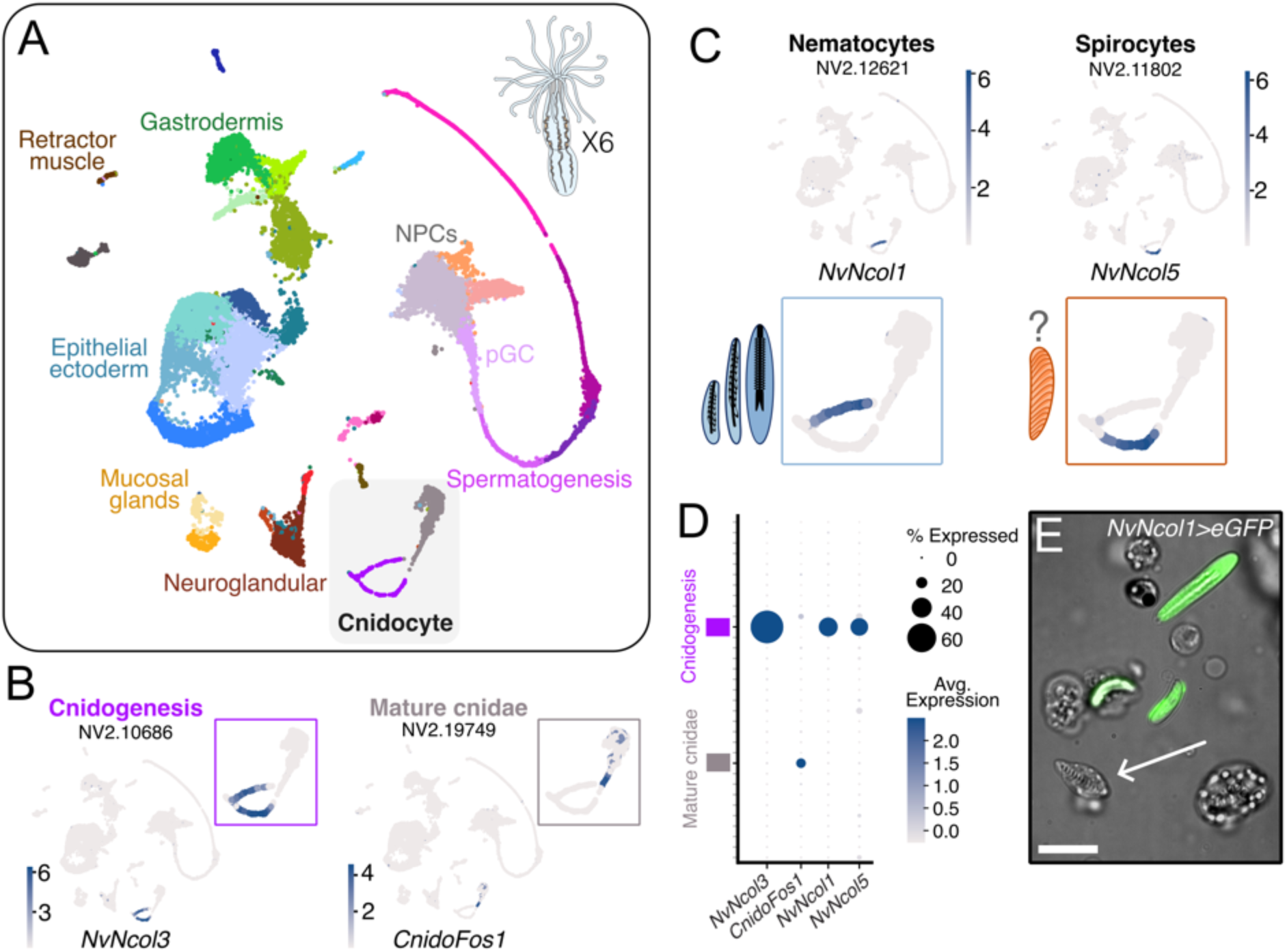
Adult *Nematostella* scRNAseq atlas indicates two distinct cnidocyte populations marked by distinct minicollagens. **A)** Whole-animal atlas from six individual adult *Nematostella*, with major cell types indicated. The putative cnidocyte clusters are highlighted in a gray square. **B)** Expression profiles from *NvNcol3*, a known cnidogenesis marker, and *CnidoFos1*, a known marker for mature cnidocytes in *Nematostella*. **C)** Expression profiles for *NvNcol1*, a known nematocyte-specific marker in developing cells, and *NvNcol5*, a putative spirocyte marker. Each display distinct expression profiles along the bifurcated cnidogenesis cluster. **D)** Dotplot of expression level for *NvNcol3*, *CnidoFos1*, *NvNcol1*, and *NvNcol5*, showing the specificity to cnidocyte-specific clusters. **E)** Dissociated cells from *NvNcol1>eGFP^cnidae^* reporter line, showing strong signal in developing and mature nematocysts but a lack of signal in the spirocyst (which remains contained in the spirocyte) (white arrow). Scale bar: 10 μm.

We designated two clusters in the whole animal dataset as either developing or mature cnidocytes based on the expression of *NvNcol3* and the mature cnidocyte marker *CnidoFos1*^34^ (**Figure 3B**). Interestingly, the developing cnidocyte cluster encompassed two distinct subclusters which converged on the mature cnidocyte cluster, a cell trajectory described in previous *Nematostella* single cell atlases^18^. One of these developing cnidocyte subclusters expressed *NvNcol1* while the other expressed *NvNcol5* (**Figure 3C, D**), suggesting a possible divergence between nematocytes and spirocytes, respectively. Indeed, we confirmed the absence of *NvNcol1>eGFP*^cnidae^ expression in spirocytes from dissociated adult animals (**Figure 3E**). We thus identified two major cnidocyte cell types in this analysis, namely *NvNcol3+/NvNcol1+* nematocytes and a distinct *NvNcol3+/NvNcol5+* cnidocyte type, which likely represents spirocytes (**Figure 3**).

To identify additional cnidocyte-specific expression patterns, we next performed subclustering analysis of the developing and mature cnidocyte clusters **(Figure 4**). This approach identified two distinct *NvNcol3+* cell populations that we hypothesize to reflect transcriptional differences between developing nematocytes and spirocytes (**Figure 4**, blue and orange clusters, and **Figure 4E**). Supporting this interpretation, *NvNcol1* and *NvNcol6* were within the top five markers for the nematocyte-associated cluster (cluster 5; **Figure 4C**). In contrast, *NvNcol5* was in the top five markers for the putative spirocyte cluster (cluster 2; **Figure 4C**). Consistent with our expectation that the constant turnover of cnidocytes ensures capture of various stages of development, both clusters shared expression of the early cnidocyte markers *NvSox2*^38^, *NvZnf845*^37^, and *NvPaxA*^36^ as well as the late-stage cnidocyte marker *CnidoFos1*^34^. This could suggest similar cellular trajectory-like patterns in both *NvNcol3+* cluster groups (**Figure 4D**). *NvSoxA*^18^, a mature cnidocyte marker, was present in a separate, larger cluster group corresponding to the putative mature cluster from the whole tissue UMAP (**Figure 4D**). Other cnidocyte-specific structural genes were expressed in both developing cnidocyte cluster groups, while the nematocyte-specific protein encoding *spinalin-like* or *v1g243188* (NV.10343)^17,35^ was expressed only in the nematocyte-associated clusters (**Figure 4E**). Notably, the previously described spirocyte marker *NvCaluF* was expressed solely in the *NvNcol3+*/*NvNcol1-*cluster (**Figure 4E**). In summary, our subclustering analysis of the whole adult dataset revealed distinct expression dynamics for clusters associated with developing nematocytes and putatively developing spirocytes.

**Figure 4:**
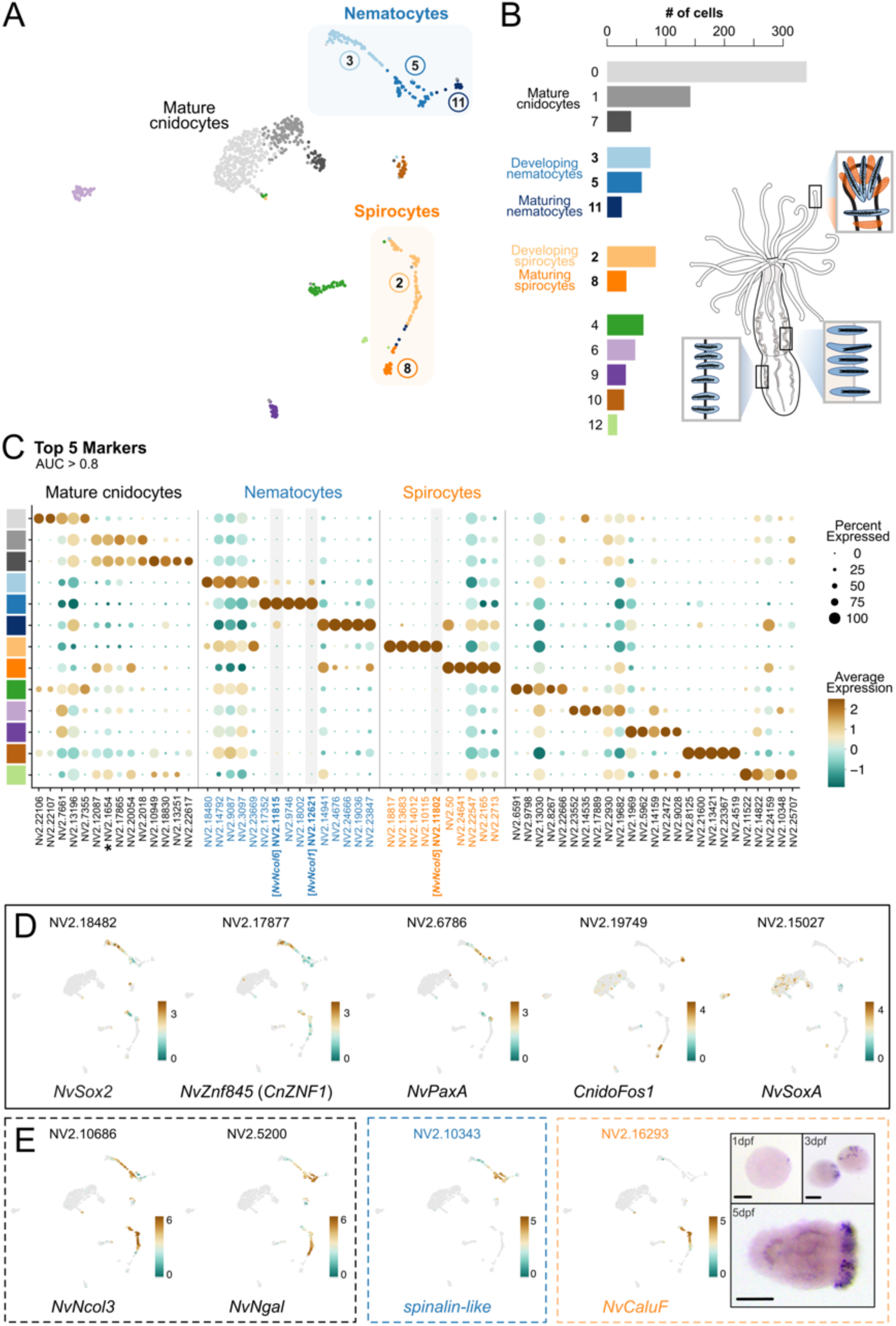
Re-clustering analysis refines the transcriptomic profiles of adult nematocytes and spirocytes. **A)** Atlas of re-clustering scRNA-seq from whole, adult dataset, with putative developing nematocytes and spirocytes indicated, in additional to putative mature cnidocyte clusters. **B)** Number of cells across each of the defined cell clusters. Schematic of adult *Nematostella* indicates the distribution of cnidae morphologies across adult tissues. **C)** Dotplot indicating top five markers for each of the stinging cell subset clusters, including the putative mature cnidocytes, nematocytes, and spirocytes. Gray bars indicate minicollagen genes that are top markers in putative developing nematocyte and spirocyte clusters indicated in A. *Indicates a marker that was a top five marker present in two different clusters within the mature cnidocyte clusters. **D)** Expression plots of known cnidogenesis and mature cnidocyte markers from previous studies. **E)** Expression plots for specific cell subtype-specific markers. Black dotted box: Known cnidocyte specific structural proteins, including *NvNcol3* and *NvNematogalectin* (*NvNgal*). Blue dotted box: Known nematocyst specific structural protein, annotated as spinalin-like (^17^ but see ^35^). Orange dotted box: spirocyte-specific marker and complimentary CISH imaging across early developmental stages (late gastrula (1dpf), early planula (3dpf), and budding planula(5dpf)), showing expression is present only in oral/developing tentacle region in later developmental stages, which is indicative of spirocyte specificity.

### NvNcol5 marks developing spirocytes

Leveraging our whole tissue scRNA-seq dataset, we next sought to characterize expression dynamics across the minicollagens. Our cnidocyte-specific scRNA-seq dataset shows distinct gene expression patterning of minicollagens between nematocytes and putative spirocytes (**Figure 5**). *NvNcol1, NvNcol4,* and *NvNcol6* were expressed within the nematocyte-specific clusters (nematogenesis), while *NvNCol5* and *NvNcol0* were expressed only within the putative spirocyte cluster (“spirogenesis”). Only *NvNcol3* was expressed within both cluster groups. Expression patterns in each of these two developing cnidocyte clusters further exhibited a degree of temporal specificity. In the case of nematogenesis, *NvNcol3* appears to be an early cnidocyte marker, given the expression of early cnidocyte transcription factors (**Figure 4D**). *NvNcol3+* expression is followed by *NvNcol1* and then *NvNcol6,* while *NvNcol4* appears to have a more constricted expression pattern that overlaps with the other three minicollagens (**Figure 5B**). In the putative spirocyte clusters, *NvNcol3* appears to be an early marker followed by *NvNcol5* and *NvNcol0*, (**Figure 5C**). The observation that *NvNcol6* and *NvNcol0* were expressed later in cnidogenesis is congruent with our HCR results, where multiple cells displayed only *NvNcol6+* or only *NvNcol0+* signal, and absence of *NvNcol3* (**Figure 2A**). We further quantified the overlap of HCR signal from *NvNcol5+* cells against the nematocyte-restricted *NvNcol1* and *NvNcol4* in multiple late-stage planulae (n=7). Little overlap of *NvNcol5* with either *NvNcol1* or *NvNcol4* was observed, but there was statistically higher overlap of *NvNcol1* and *NvNcol4* (**Figure 5D**). This provides strong evidence that: 1) *NvNcol4* is a nematocyte-specific marker that overlaps with a subset of *NvNcol1+* cells, reaffirming the restricted expression observed in our scRNA-seq; and 2) *NvNcol5* marks a distinct cnidocyte population, likely developing spirocytes.

**Figure 5:**
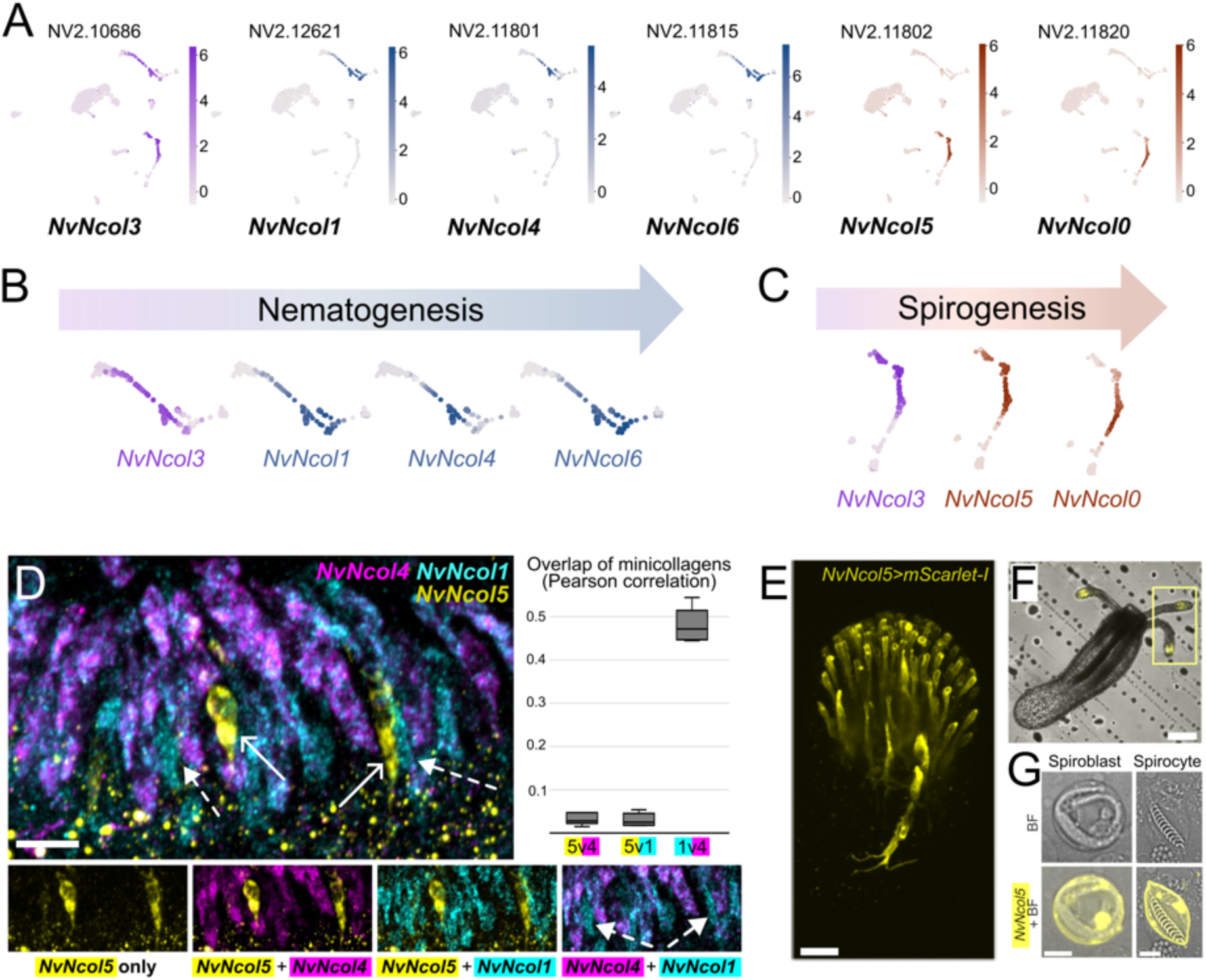
Minicollagen expression dynamics in developing nematocytes and spirocytes from adult *Nematostella*. **A)** Expression plots from each of the six minicollagens used for HCR experiments, as shown in Figure 2. Purple indicates the universal cnidocyte marker *NvNcol3*, blue indicates nematocyte-specificity, and orange-red indicates spirocyte-specificity. **B) and C)** Close-up of the minicollagen expression patterns in developing cnidocyte clusters shown in **A)**, indicating both cluster-specificity and putative temporal specificity across **B)** nematogensis and **C)** spirogenesis. **D)** Quantitative overlap analysis of *NvNcol5*, *NvNcol1*, and *NvNcol4* from HCR experiments on 4dpf animals. Image is representative of the budding tentacle. White solid arrows indicated *NvNcol5+* cells with no overlap of *NvNcol1* or *NvNcol4*. While majority of cells are *NvNcol1+/NvNcol4+*, white dotted arrows indicate cells that are solely *NvNcol1+*. A Pearson correlation quantification that indicates the overlap of each of the three genes is shown. **E)** Tentacle tip of F1 polyp from a *NvNcol5>mScarlet-*I transgenic line. **F)** mScarlet-I signal at tentacle tips of F1 *NvNcol5>mScarlet-*I primary polyp. **G)** Dissociated cells from *NvNcol5>mScarlet-*I F2 adult animals showcasing mScarlet-I signal in a developing spirocyte (spiroblast) and nearly or recently mature spirocyte. Scale bars: **D, G)** 5 μm, **E)** 10 μm, **F)** 100 μm.

To confirm that *NvNcol5+* cells are indeed spirocytes, we created a transgenic reporter line using 966 bp of the upstream regulatory region of *NvNcol5* to drive mScarlet-I *in vivo*, called *NvNcol5>mScarlet-I*. In both F1 and F2 heterozygous primary polyps mScarlet-I expression was only observed at the tentacle tips, matching the observed patterns in our HCR experiments (**Figure 5E,F**). As these animals matured and additional tentacles developed, mScarlet-I+ cells expanded across the tentacle tissue from the tip to the base. We dissociated adult tissues from *NvNcol5>mScarlet-I* and observed mScarlet-I in developing spirocytes (i.e. spiroblasts) and recently matured spirocytes (**Figure 5G**). In contrast, reporter expression was absent from developing and mature nematocytes. Thus, through multiplex HCR experimentation and signal overlap quantification, adult scRNA-seq analysis and transgenesis, we confirm that *NvNcol5* is a faithful marker for developing spirocytes.

### RNA-FACS-sequencing reveals distinct transcriptional profiles during development of nematocytes and spirocytes

Given the robust expression of minicollagens in early development and their cnidocyte subtype-specific expression patterns inferred from scRNA-seq (**Figures 2 and 5**), we next sought to obtain deeper transcriptional insights using RNA-FACS-seq^51–53^, a modification of Probe-seq^54^ (**Figure 6A**). Briefly, approximately 50 *wild-type* adult animals from the same spawning group were collected, raised to adulthood, and subsequently fixed and dissociated into single cells. The resulting cell suspension was split into six samples (plus two gating control samples). Each sample was fluorescently labeled for one of the six minicollagens using HCR and then sorted using FACS. Approximately 1,500 cells/replicate were collected from each labeled cell population for bulk RNA-seq analysis (**Supplemental Data S3**). Differentially expressed genes (DEGs) were determined using edgeR, and the results are summarized in **Supplemental Data S3** with annotations using DIAMOND *blastp* searches against UniProt SwissProt, UniProt TREMBL, and the NCBI nr protein database as well as the top domain matches against PFAM. All together, we produced replicate bulk RNA-seq datasets for each of the six minicollagen-expressing cell populations that were suitable for differential expression analyses.

**Figure 6:**
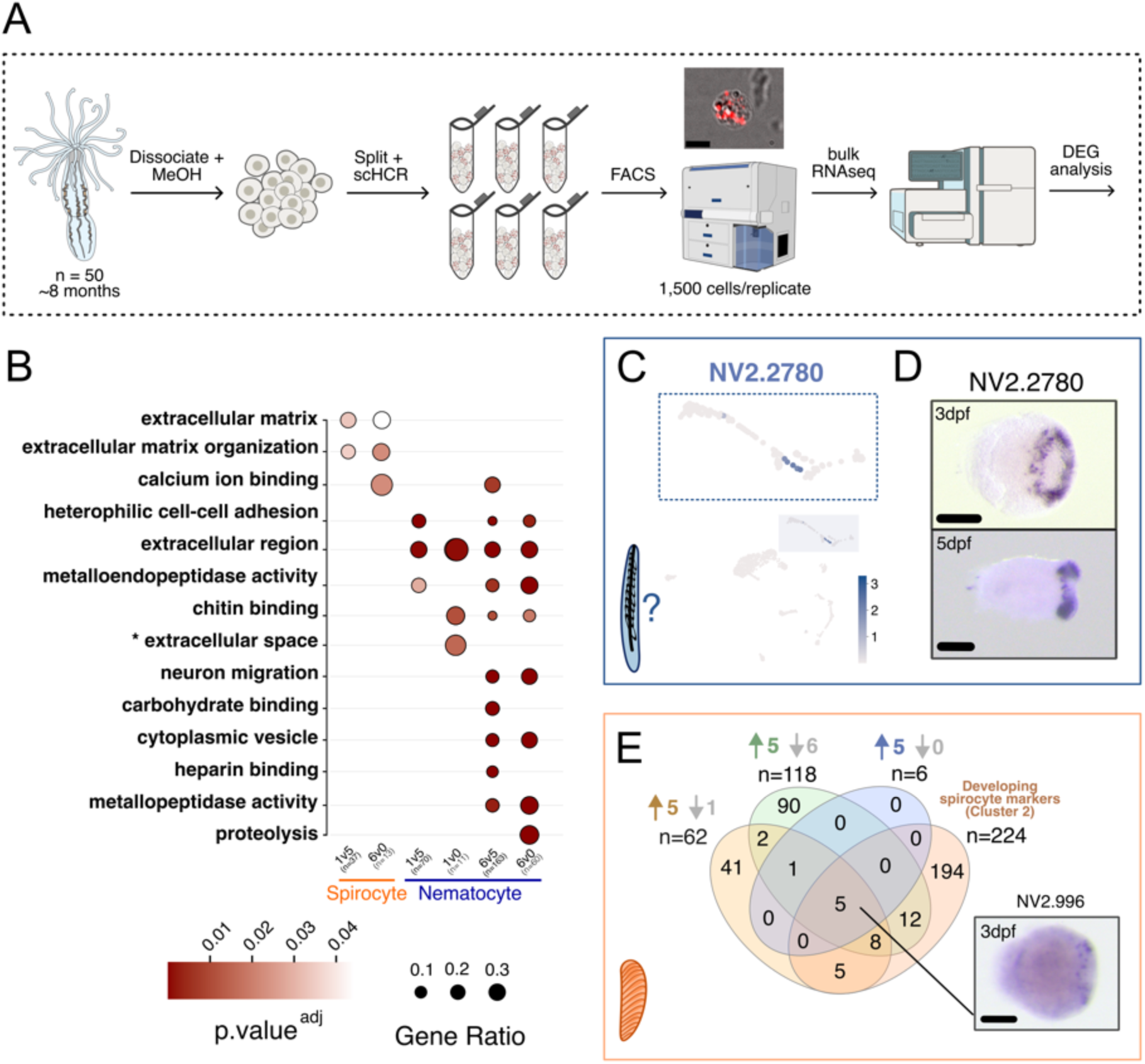
RNA-FACS-seq analysis of six HCR-labeled adult *Nematostella* cell populations. **A)** Schematic of RNA-FACS-sequencing pipeline used in this experiment. Image above FACS machine indicates a cell with positive HCR signal, post sorting. Multiple images used from NIH Bioart Archive. **B)** GO enrichment analysis from DEG genes determined using edgeR through comparisons of spirocyte-associated cell populations (*NvNcol5* and *NvNcol0*) against nematocyte-associated cell populations (*NvNcol1* and *NvNcol6*). Color denotes significance after BH correction (adjusted p-value); dot size indicates the number of genes enriched across all total genes analyzed within each sample, shown as n = # below each comparison. **C)** Expression plot of NV2.2780, which was found to be an up-regulated gene in *NvNcol6* population but not identified as a scRNA-seq marker gene. **D)** CISH pattern of NV2.2780 in 3dpf and 5dpf animals, showcasing unusual expression in early development with majority of signal only at the oral opening and developing tentacles. **E)** Venn diagrams indicating overlap of DEGs from edgeR analysis up-regulated in *NvNcol5* cell population compared to *NvNcol1*, *NvNcol6*, and *NvNcol0* as well as the markers for cluster 2 in the cnidocyte-specific cell scRNA-seq, in which *NvNcol5* was identified within the top five markers. Venn diagram modified from InteractiVenn^79^. CISH image of NV2.996 expression in a 3dpf animal. Scale bar: **A)** 5 μm, **D,E)** 100 μm.

Using edgeR analysis, we identified several hundred genes that were differentially expressed between NvNcol1/6-expressing nematocytes and NvNcol0/5-expressing spirocytes (**Supplemental Data S3**). We consistently found a higher number of up-regulated DEGs in the cell populations associated with nematocytes compared to spirocytes, including previously described venom toxins (**Table 1**). This matches the general expectations that: 1) Spirocysts are structurally less complex than the morphologically diverse nematocysts and thus express fewer structural genes; and 2) Spirocytes do not synthesize or deploy venoms.

**Table 1:** Expression patterns of known *Nematostella* venom toxins in this study.

| GENE | NV2 | scRNA-seq (cnidocyte subcluster) |  | RNA-FACS-seq |  |  | REF |
| --- | --- | --- | --- | --- | --- | --- | --- |
|  |  | Present in Nematocyte (cluster #) | Present in Spirocyte (cluster #) | DEG (Y/N)? | CELL TYPE | TEST (bold = nem) |  |
| **NEP3** | NV2.20491 | Y (5,11) | N | Y | NEM | **1v5**<br>**6v5** | Moran *et al.* <sup>82</sup> |
| **NEP3-like** | NV2.20489 | Y (3,5,11) | N | Y | NEM | **1v5**<br>**6v5**<br>**6v0** | Moran *et al.* <sup>82</sup> |
| **NEP4** | NV2.20021 | Y (3,5) | N | Y | NEM | **6v5** | Moran *et al.* <sup>82</sup> |
| **NEP6** | NV2.18306 | Y (5) | N | Y | NEM | **1v5**<br>**1v0**<br>**6v5**<br>**6v0** | Moran *et al.* <sup>82</sup> |
| **NEP8** | NV2.19484 | Y (5) | N | Y | NEM | **6v5** | Moran *et al.* <sup>82</sup> |
| **NEP8-like** | NV2.20023 | Y (3,5) | N | Y | NEM | **1v5**<br>**6v5**<br>**6v0** | Sachkova *et al.* <sup>83</sup> |
| **NEP14** | NV2.24493 | Y (5) | N | Y | NEM | **1v5**<br>**6v5**<br>**1v5** | Moran *et al.* <sup>82</sup> |
| **NEP16** | NV2.12063 | Y (3,5) | N | Y | NEM | **6v5**<br>**6v0** | Moran *et al.* <sup>82</sup> |

Using a GO-enrichment analysis with *clusterProfiler* (function = enricher) and custom annotation of DEGs, we found relatively few GO terms significantly enriched in spirocyte-associated DEGs but multiple enriched GO terms associated with nematocyte DEGs (**Figure 6B**). Within up-regulated spirocyte DEGs (i.e. *NvNcol5* and *NvNcol0* cell populations), enriched GO terms included extracellular matrix (GO:0031012), extracellular matrix organization (GO:0030198), and calcium ion binding (GO:0005509), which may be involved in the assembly and discharge dynamics. For DEGs associated with nematocytes (i.e. *NvNcol1* and *NvNcol6* cell populations), enriched GO terms included heterophilic cell-cell adhesion (GO:0007157), extracellular region (GO:0005576), chitin binding (GO:0008160), and metalloendopeptidase activity (GO: 0004222), which may be involved in ensuring proper assembly of the subcellular features of the nematocyst organelle (e.i. barbs, spines) as well as toxin processing. Interestingly, GO terms enriched specifically in the *NvNcol6+* DEGs (i.e. not identified as up-regulated in *NvNcol1*), such as neuron migration (GO:0001764), cytoplasmic vesicle (GO:0031410), heparin binding (GO:0008210), metallopeptidase activity (GO:0008237), and proteolysis (GO:0006508). Given that *NvNcol6* appears to peak in expression later in nematogenesis based on both HCR experiments and scRNA-seq, these could be important genes involved in the maturation and migration of nematocytes. Altogether, our RNA-FACS-seq approach identified numerous genes that differ between these two subtypes with putative roles in organelle assembly, structure, and toxin production and maintenance.

To develop a set of nematocyte marker genes, we cross referenced nematocyte-associated DEGs from our RNA-FACS-seq with those identified by scRNA-seq (**Figure 6**). We first pooled up-regulated DEGs in *NvNcol1*-labeled cell populations labeled compared to 1) *NvNcol5*, 2) *NvNcol0*, and 3) *NvNcol6*, which totaled 188 unique genes. Similarly, we pooled the DEGs up-regulated in *NvNcol6+* cells against 1) *NvNcol5*, 2) *NvNcol0*, and 3) *NvNcol1*, which totaled 366 unique genes (**Supplemental Fig. S11**). We compared these two pools of up-regulated nematocyte-associated DEGs with marker genes identified from cluster 5 in our cnidocyte subcluster atlas (n = 419), where both *NvNcol1* and *NvNcol6* were in the top five markers (**Figure 4C**). In both comparisons, over half of the DEGs identified in the RNA-FACS-seq dataset overlapped with genes in the cluster 5 marker table, including respective minicollagens (**Supplemental Fig. S11)**. However, genes identified in the DEG analyses but not in the marker gene analysis of our cnidocyte-subset scRNA-seq may still have nematocyte-specific expression patterns. For example, *Hemicentin-1* (NV2.2780) was up-regulated in the *NvNcol6*+ cell populations when compared to either *NvNcol5* or *NvNcol0* and displayed relatively high expression in a small number of cells in the scRNA-seq dataset, but was not identified as a marker of this cluster (**Figure 6C**). To validate *Hemicentin-1* expression *in vivo*, we performed CISH and we observed strong signal of in the oral end and developing tentacle region of late-stage planulae (**Figure 6D**), indicative of developing cnidocytes.

To identify spirocyte-specific genes shared between our RNA-FACS-seq and scRNA-seq analyses, we compared up-regulated DEGs in the *NvNcol5+* cell population against markers identified in cluster 2 of the cnidocyte-subset (n = 224), given cluster 2 included *NvNcol5* as a top marker (**Figure 4C**). We found there was less relative overlap between spirocyte-specific genes across these two datasets when compared to the nematocyte markers noted above (**Figure 6E**). We did identify five genes that were shared between all of our DEG analyses and scRNA-seq, including a gene annotated as *Coagulation factor VIII* (NV2.996; **Figure 6E**).

*Coagulation factor VIII* was sparsely expressed at the oral pole of planula larvae using CISH (**Figure 6E**), which may correspond to developing spirocytes based on our previous CISH labeling of *NvCaluF* (**Figure 4E**). This suggests that our RNA-FACS-seq method successfully identified spirocyte-associated candidates for future study, though the variability between this dataset and our scRNA-seq marker analysis will require additional validation. Overall, these comparisons show that our RNA-FACS-seq strategy acquired high-quality transcriptional data from low cell count, HCR-labeled cell populations, which we believe is the first application of this methodology in *Nematostella*. Our DEG results complement our findings from our scRNA-seq for both nematocyte-specific and spricoyte-specific genes but, importantly, identified cell type-specific candidates that were masked in our scRNA-seq experiment.

## DISCUSSION

Given their relative ease of observation under light microscopy and exquisite morphological variations, cnidocytes have long fascinated biologists. Surprisingly, however, we still know very little about the molecular mechanisms that drive the morphogenesis of distinct cnidae types. Here, through the integration of multiplex HCR RNA-FISH experiments, adult tissue scRNA-seq and production of two transgenic lines, we analyzed the expression dynamics of multiple members of the minicollagen protein family in *Nematostella* cnidocytes. This led to the identification of two minicollagens, *NvNcol5* and *NvNcol0,* as robust markers for the spirocyte lineage. We also found that the minicollagens display nuanced but distinguishable temporal expression patterns across nematogenesis and spirogenesis. Aggregating this data, we propose the following transcriptional sequence: 1) *NvNcol3* is an early and ubiquitous marker for both nematocytes and spirocytes; 2) Nematocytes subsequently express *NvNcol1*, *NvNcol4*, and *NvNcol6*, where *NvNcol1* and *NvNcol4* are early-to-mid-nematogenesis markers, and *NvNcol6* is a mid-to-late nematogenesis marker; and 3) *NvNcol5* is an early-mid marker of spirogenesis marker followed by expression of *NvNcol0* (**Figure 7**). This new framework provides the foundation for future molecular, functional, and evolutionary studies of these two major cnidocyte types.

**Figure 7:**
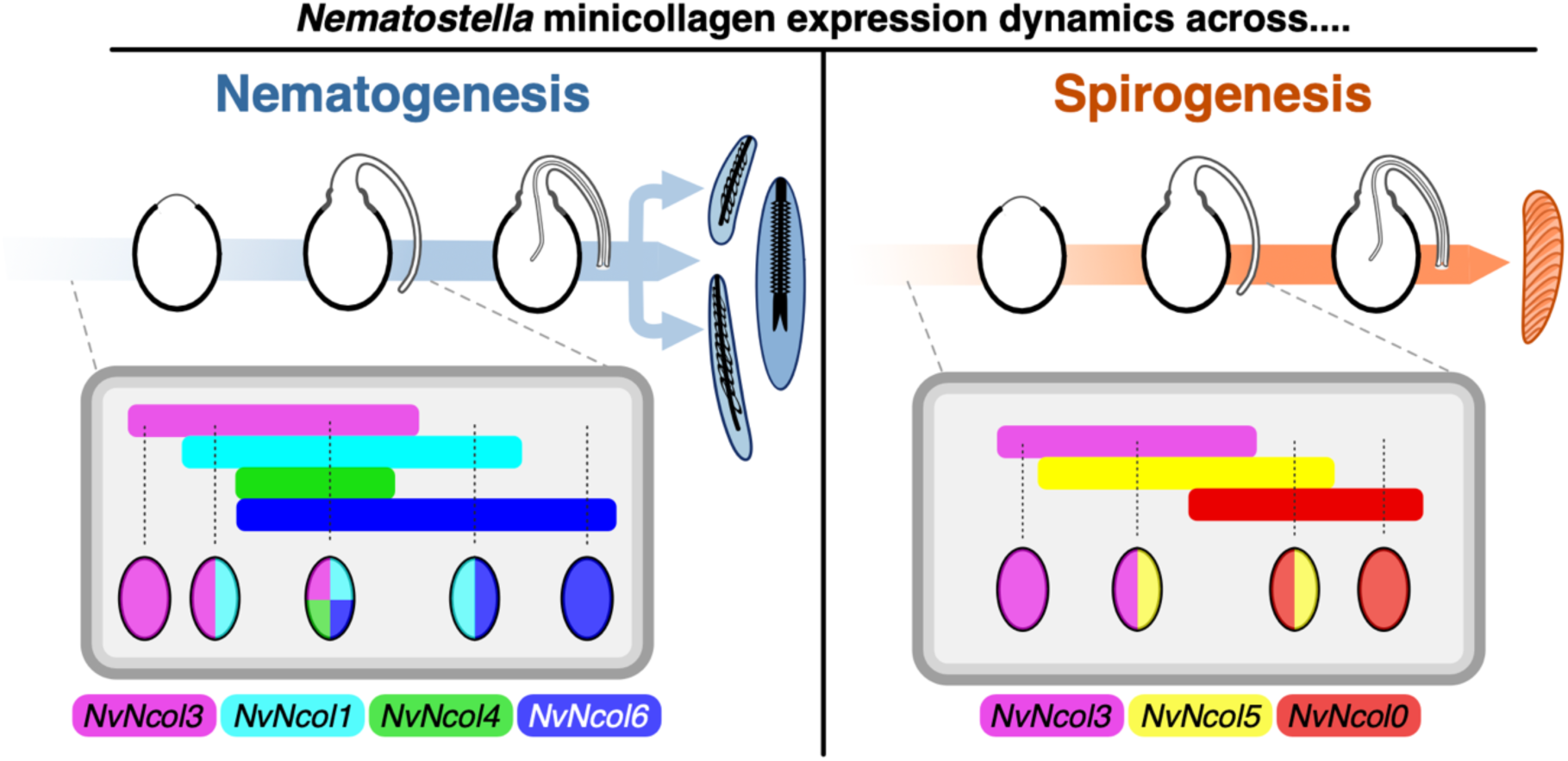
Transcriptional program of nematogenesis and spirogenesis summarized by minicollagen expression, as shown in this study. Schematic of cnidogenesis is modified from Holstein 1981^80^ and Skaer 1973^81^ depicting capsule formation, tubule formation, and inversion of the tubule into the capsule with arrows indicating maturation of cnidae, either nematocysts (blue) or spirocysts (orange). Boxes indicate relative overlap of expression for each minicollagen gene during cnidogenesis. Ovals indicate specific predicted cell populations and respective gene expression patterns. For example, populations of *NvNcol3+* only cells are expected to populate during early nematogensis and spirogenesis. In the next stages of nematogenesis, populations of *NvNcol3+/NvNcol1+* (purple/blue) indicates cell populations at early nematogenesis, shown by the overlap of the representative purple and blue bars.

To extend our understanding of cnidocyte specification beyond minicollagen dynamics, this study reports the first bulk transcriptomic profiling of a fixed, FACS-sorted cell population from *Nematostella*. This analysis yielded hundreds of DEGs, finding that significantly more genes were up-regulated in nematocytes than in spirocytes. Genes upregulated in nematocytes included several encoding venom proteins, which validates a long-standing prediction that spirocysts do not deploy venom toxins, as least in this species (**Table 1**). GO-term analysis of these DEGs further highlights multiple functionally interesting candidates for downstream analysis, in particular for the later developing *NvNcol6+* nematocyte population. For example, we found that *Hemicentin-1* (NV2.2780) was up-regulated in *NvNcol6+* cell populations but was not identified as a marker in our scRNA-seq analysis. We thus directly analyzed *Hemicentin-1* expression and found it was restricted to the developing tentacles, suggesting it may be specifically expressed in large isorhizae nematocytes (**Figure 6D**). Interestingly, *Hemicentin-1* was identified as a cluster marker in a previous scRNA-seq dataset annotated as “nem.2”, which was hypothesized to be a specific nematocyte-type, such as large isorhizae^18^. This showcases the utility of our RNA-FACS-seq methodology for identifying highly cell type-specific genes that may be masked in conventional scRNA-seq analyses. Furthermore, our whole adult scRNAseq and RNA-FACS-seq datasets provides a wealth of potential cnidogenesis candidates for further interrogation into nematocyte and spirocyte biology.

Cnidocytes are important examples of cellular novelties, and *Nematostella* has been become a central model to interrogate their specification and morphogenesis. While previous work has typically focused on the broader cnidocyte lineage, *Nematostella* displays distinct cnidocyte subtypes and thus presents an opportunity to determine the molecular mechanisms underlying lineage diversification and cell type evolution. In this study, we established a transcriptional framework to distinguish *Nematostella* nematocyte and spirocyte development using minicollagen expression. In addition to the ability to dissect functional properties of these two subtypes, such as differing structural components or the gain/loss of venom toxins, our findings will also facilitate future functional studies on the gene regulatory networks that distinguish them. Our transcriptional framework for nematogenesis and spirogenesis thus advances the molecular tools, gene expression datasets, and foundational biological insights that showcase *Nematostella* cnidocytes as a highly compelling model within the field of evolutionary cell biology^9,34,37,41,42^ and functional genomics^35,36,38–40^.

On a technical level, this work demonstrates the utility of a bulk RNA-sequencing approach to characterize rare or delicate cell populations though the application of an optimized RNA-FACS-seq approach. While our scRNA-seq analysis provided a holistic overview of the cell types present in adult tissues, the dataset only sparsely represented specific, rare cell types. Our FACS-based approach to sorting fixed, HCR RNA FISH-labeled cells offers several distinct advantages. First, RNA-FACS-seq uses fixed cell samples, which enables the collection of rare or fragile cell populations in a specific tissue or life stage of interest. Second, RNA-FACS-seq takes advantage of *wild-type* individuals and is not limited to laboratory models where genomic-manipulation is feasible. Third, while the use of FACS-enriched cells from transgenic lines has been employed by other studies for characterization of cnidocytes^36,42^, including this study, RNA-FACS-seq isolates a cell population at the precise time of target RNA expression. In contrast, fluorophore expressed in transgenic reporter lines require time to mature, distorting temporal dynamics. Furthermore, reporter lines marking fragile cell types may also remain challenging to isolate at sufficient quality for scRNA-seq as compared to a fixed cell method. Lastly, while an alternative strategy could use antibodies to label specific cell populations, this would again mark cell populations at the time of protein maturation and not peak RNA expression. These differences aside, for major cell types we found reasonable repeatability of the gene expression patterns observed across experimental modalities. We believe that the RNA-FACS-seq approach reported here showcases a practical experimental strategy for those working with non-traditional model organisms.

## METHODS

### Animal Husbandry and Spawning

Animals were maintained in Pyrex dishes in freshly mixed 12ppt artificial sea water (ASW) (Instant Ocean). Active spawning colonies were under the care of the Stowers Invertebrate Team and maintained at 16C on a 12:12 light-dark cycle, fed 2-5X a week on *Artemia fransiscana* nauplii, and spawning and fertilization conducted as previously described^55–57^. All experimental animals were maintained at ambient room temperature (RT) (∼20-22C) and fed 1-2X per week with *Artemia* nauplii and water changed the following day, unless otherwise specified. All animals used in HCR experiments were *wild-type* and raised at RT unless otherwise specified.

Adult animals raised for HCR RNA-FACS-seq experiments were fertilized on the same day with clonal spawning colonies and cultured in a single dish at RT with ambient light. Animals were initially fed 3X per week with *Artemia* and then transitioned to 1X every two weeks after ∼3 months, which water changes occurring the day after a feeding. The reduced feeding schedule ensured animals remained at a manageable size for whole tissue dissociations, as larger animals were more difficult to dissociate. Animals used for planulae-specific HCR RNA-FACS-seq were spawned separately and maintained in the same dish with ambient light to 4dpf at RT.

### Generation of Transgenic Lines

The *NvNcol1>eGFP^cnidae^*, *NvNcol1>eGFP*, and *NvNcol5>mScarlet-I* transgenic lines were generated by meganuclease as previously described^58,35^. For the *NvNcol1>eGFP^cnidae^* line, the first and second exon, first intron, and a portion of the second intronic region were included. Primers used for amplification of promoter regions from the NV2 genome assembly, available at Stowers Institute for Medical Research (https://simrbase.stowers.org/), are provided in the **Supplemental Data S1**. Microinjection was carried out in unfertilized eggs using a pulled glass capillary with the FemtoJet 4x system (Eppendorf). A single F0 animal from each line was used to produce F1 animals, and a single F1 individual was used to establish an F2 generation. Unless otherwise specified, F2 or subsequent generations were used for imaging.

### Whole Tissue Hybridization Chain Reaction (HCR) RNA-FISH

The protocol for whole tissue HCR RNA-FISH was modified from Molecular Instruments v3 protocol and previous protocols for planaria^52^ (see **Supplementary File S1**). Probes for each gene were ordered from Molecular Instruments; gene models are presented in **Supplemental Data S1**. The following probe set amounts were used in 200 μl total probe solution steps: *NvNcol3*: 5 μl, *NvNcol1*: 5 μl, *NvNcol4*: 5 μl, *NvNcol6*: 5 μl, *NvNcol5*: 5 μl, and *NvNcol0*: 10 μl. For 3-color HCR experiments, AlexaFluor 488, 546, and 647 nm amplifiers were used for imaging on an Andor DragonFly 200; for 6-color HCR experiments AlexaFluor 488, 546, 555, 594, 647, and 700 nm amplifiers were imaged on a Leica SP8 Confocal STED (see below).

### Colorimetric *in situ* Hybridization (CISH)

CISH for whole tissue followed the standard protocol as previously described, including probe development^59^. Primers used for probe synthesis are listed in **Supplemental Data S1**. Probes were synthesized using a MEGAscript T7 Transcription Kit (Invitrogen, #AM1334) from templates that were either after transformation of the amplified region into a Zero Blunt TOPO pc4 plasmid (Invitrogen, #450031) or directly from the PCR product of a cDNA template, where T7 region was included in the reverse primer.

### Imaging, Analysis, and Quantification

Most images were acquired on an Andor DragonFly 200 Confocal Microscope (Oxford Instruments). For live images, whole animals were immobilized using several drops of 7% MgCl2 • 6H20 in 12ppt ASW until little movement was observed. For fixed images, tissue was mounted in a mounting solution of 90% glycerol with 1μl of DAPI (10mg/ml) in 10mL of mounting solution. For 6-color HCR samples, images were taken using a SP8 Confocal STED with Falcon Lifetime (Leica) with standard fluorescence channels. However, amplifiers for AlexaFluor 546 and AlexaFluor 555 were separated using a custom python script to unmix the dyes with phasor unmixing. Amplifier-only controls are shown in **Supplemental Fig. S2**. Images were processed using Fiji (ImageJ2, v2.14.0/1.54f, build: c89e8500e4)^60,61^.

For quantification of HCR signal from *NvNcol1*, *NvNcol4*, and *NvNcol5,* images from six planulae were segmented in 3D using SAM2^62^ after drawing an initial prompt around one slice using *napari*^63^. This segmentation was used to mask out areas on the periphery of the image, as non-specific staining on the surface of the animal was high. Depth attenuation of fluorescence signal was corrected by dividing by a decaying exponential on Z. Images were blurred and masks of high intensity regions of *NvNcol1*, *NvNcol4*, and *NvNcol5* were generated based on otsu or manual thresholding. Pearson’s correlation coefficient was calculated for each combination of the three masks using scikit-image^64^. All analysis was completed in Python.

### Whole-animal Single-cell RNA Sequencing (scRNA-seq) of Adult *Nematostella*

#### Dissociation of Adult Tissue

Dissociations for adult single cell experiments were conducted as outlined by Steger *et al.* ^18^. Three 16-tentacle staged male *Nematostella* from the *NvNematogalectin>mScarlet-I* line^35^ were starved 3 days and cleaned from debris prior to dissociation. Each animal was visualized to ensure fluorescent signal. Animals were collected into a 5ml tube pretreated with PBS+ 0.1% Tween (PTw) and washed 3X with ASW. Animals were briefly washed in two bowls of fresh ASW and allowed ∼20-30min to relax in the tube. As much ASW as feasible was removed and replaced with 10X TrypLE (Gibco, #A1217701), and dissociation was conducted with gentle constant rocking at RT with occasional (∼15 min) manual pipetting with a clean glass pipette.

After 2.5 hours, the reaction was quenched with 1% BSA/PTw and filtered using sterile 70μm Flowmi Tip Strainers (Sp-Bel-Art, #136800070). Cells were pelleted at 4C, 300rcf for 5 min and washed twice with 1% BSA/PTw. Cells were resuspended in 1mL 1% BSA/PTw and maintained on ice prior to FACS. A similar dissociation with three WT male individuals at the 16-tentacle stage was used for an additional scRNA-seq experiment. Dissociation was carried out as above and cells transported for cell viability assays.

#### Fluorescence-Activated Cell Sorting (FACS) of NvNematogalectin>mScarlet-I Transgenic Line

For adult samples for scRNA-seq from the *NvNematogalectin>mScarlet-I* line, cells were briefly incubated with 1:1000 DAPI (1mg/mL) and 1:200 Draq5 (5 mM) and sorted based on viability. FACS was conducted at the Stowers Institute using a six-laser configured FACSymphony S6 Cell Sorter (BD Biosciences, USA) a 100μm nozzle and 1X sheath (PBS); the 355, 561, and 637 nm lasers were used for detection of DAPI, mScarlet, and Draq5, respectively. Samples were maintained at 4C for the duration of the sort. Data was analyzed using FACSDiva v9.1.2 (BD Biosciences, USA). Cells were sorted into a 1.7ml tube pretreated with 1% BSA/PBS, pelleted, and transported on ice for scRNA-seq library preparation.

#### Library Preparation and Sequencing - 10X Chromium

The concentration and viability of cells for both sorted and WT cell solutions was assessed using a Luna-FL cell counter (Logos Biosystems). Cells with a viability of 90% or higher were loaded onto a Chromium X Series instrument (10x Genomics) and libraries were prepared using the Chromium Next GEM Single Cell 3 ‘Reagent Kit v3.1 (10x Genomics) according to manufacturer’s instructions. Generated cDNA and short fragment libraries underwent quality and quantity assessment using a 2100 Bioanalyzer (Agilent Technologies) and Qubit Fluorometer (Thermo Fisher Scientific). Libraries were pooled and converted for processing on the G4 sequencer (Singular Genomics) with the SG Library Compatibility Kit, following the “Adapting Libraries for the G4 – Retaining Original Indices” protocol. The converted pool was sequenced on an F3 flow cell (Cat. #700125) on the G4 instrument with the PP1 and PP2 custom index primers included in the SG Library Compatibility Kit (Cat. #700141), using instrument software versions current at the time of processing to target a minimum average of 25,000 reads per cell with paired read lengths of 28*10*10*90 bp.

#### Data Processing and Analysis

scRNA-seq data from each of the three samples (two FACS-enriched replicates and one WT sample) were first processed using 10x Genomics Cell Ranger software (v4.0.0) (**Supplemental Data S2**). Raw reads were demultiplexed into FASTQ format using cellranger *mkfastq*. Alignment to the *Nematostella* NV2 genome assembly via the SIMR database (see **Data Availability**), filtering, barcode counting, and UMI counting was processed using the cellranger count pipeline. Featured gene matrices were then processed and analyzed using the Seurat package (v4)^65^ in R (v4.3.1)^66^. The analysis followed the standard Seurat workflow for quality control, normalization, unsupervised clustering (resolution 0.8), and UMAP visualization (**Supplemental Fig. S4**). Marker tables were produced using *FindAllMarkers* with only.pos = “TRUE” and test= “MAST” (**Supplemental Data S2**). For the cnidocyte-specific dataset, clusters 9 and 22 from the combined dataset were processed as above with clustering performed at resolution = 0.5 (**Supplemental Fig. S7**). Marker tables were produced using the *FindAllMarkers* with test= “roc” (**Supplemental Data S2**). Gene expression was visualized from both datasets using the DotPlot function. Clusters were manually annotated based on previously established markers and marker genes defined by Cole *et al.* ^19^ (**Supplemental Data S2**). A clustered dot plot for the top five markers of each cluster in the cnidocyte subset was created using the *Clustered_DotPlot* function of the ssCustomize package (v3.0.1)^66^ (**Supplemental Fig. S8**). An interactive Shiny app was created for data visualization purposes for both the whole tissue, combined and developing and mature cnidocyte-subset UMAP reduction plots (see **Data Availability**).

### RNA-FACS-seq

#### Dissociation of Adults and Planulae

Approximately 50 adult animals (33 weeks post-fertilization) ranging between the 12-16 tentacles stage were starved for 3 days and manually cleaned from debris prior to dissociation. Approximately 25 animals in two batches were moved to a clean petri dish with ASW and individuals swiftly cut using a sterile disposable scalpel (Sklar, size 10) and immediately transferred to a 5ml RNase-free tube using a plastic transfer pipette. Excess water was removed and 2ml of Accutase cell dissociation reagent (StemPro, #A1110501) was added.

Tubes were gently rocked at 37C for 25 minutes, removed and gently pipetted up and down for ∼5 min with a sterile 21.5 gauge needle (BD Precision Glide, #305167) attached to a 1ml syringe (Norm-Jet, 4010.200V0), and then moved back to 37C for 20 minutes. After incubation, gentle pipetting for ∼5 min with a clean 26.5 gauge needle (BD Precision Glide, #305111) attached to 1ml syringe was used to bring the remaining tissue into suspension. A similar protocol was used for dissociation of ∼100 planulae at 4dpf, without the additional cutting of tissues prior to adding the dissociation solution.

#### Fixation of Cell Solutions for HCR

The cell fixation protocol is modified from Gutiérrez-Franco *et al.* ^68^, and used for both adult and planula cell suspensions. To quench the dissociation solution, 1ml of Dulbecco’s Modified Eagle Medium (DMEM; Gibco, #11995-065) supplemented with 1% GlutaMAX (Gibco, #35050-061) and 15μL of DNAse I (New England Biolabs, #M0303) were added. Samples were divided into 1.7ml tubes and spun down at 4C, ∼500rcf (2000 rpm, Microfuge 22R centrifuge). The supernatant was removed and cell pellet resuspended in 0.4mg/mL BSA in Dulbecco’s Phosphate-Buffered Saline (DPBS). The samples were centrifuged and resuspended twice as above. After the third spin, cells were resuspended in 200μl ice-cold DPBS and 800uL of 100% ice-cold methanol was added drop-wise on ice. Afterwards, samples were gently mixed and left on ice for 30 min. Cells were stored at −20C overnight.

#### HCR RNA-FISH of Cell Suspensions

Cell suspensions were resuspended in a stepwise 50/75/100% gradient in 0.01% BSA, 0.2 U/μL RNAse inhibitor (Promega, #N2111), and 1mM DTT (Promega, #P117C). The cell suspension protocol for HCR was modified from the mammalian cell in suspension protocol provided by Molecular Instruments (v9, 2023-02-13; Molecular Instruments). After resuspension, cells were washed twice in PTw and the final pellet resuspended in 400ul of Probe Hybridization Buffer for Cells in Suspension (Molecular Instruments), pre-hybridized for 30min at 37C, and brought to a final volume of 500μl with the addition of 100 μl probe solution (containing 2μL of 1uM probe stock/each probe set for each experiment). Samples were incubated at 37C overnight. After incubation cells were pelleted and the probe solution was replaced with Probe Wash Solution from Molecular Instruments. The pellet was washed twice with Probe Wash Solution, pelleted again and then the solution was replaced with 500μl 5X SSC + 0.1% Tween (SSCT) and incubated for 5 min at RT. SSCT was removed and replaced with 150μl Amplification Buffer from Molecular Instruments, and allowed to incubate for 30min at RT. For hairpins preparation, 5μL of 3μM hairpin pairs (h1 and h2) were heated to 95C for 90 seconds and cooled to RT in the dark. Hairpins were combined with Amplification Buffer to a final solution of 100μl, which was added to the HCR sample (final solution volume = 250μl). Samples were incubated overnight in the dark at RT. For RNA-FACS-seq experiments, all samples were hybridized with AlexaFluor 647 amplifiers. Prior to downstream analysis, samples were pelleted and hairpin solution removed, washed 6X with SSCT, and resuspended in PBS.

#### FACS of HCR-labeled Cell Suspensions

For RNA-FACS-seq experiments, samples were filtered through 70μm Flowmi Tip Strainers pre-moistened with 1X PBS just prior to sorting and briefly incubated with 1:1000 DAPI (1mg/mL). FACS was conducted using a six-laser configured FACSymphony S6 Cell Sorter equipped with a 70μm nozzle and 1X sheath (PBS) using 355 nm and 637 nm lasers to detect DAPI and AF647, respectively, and analyzed as above using FACSDiva v9.1.2 (representative images on **Supplemental Fig. S9**). All cell populations were labeled with the same secondary such that the same gating strategy could be used across all samples, to reduce cell bias. For adult samples, ten replicates of 1,500 cells were sorted into chilled 96-well plates containing 7μl 1X lysis reaction buffer from the SMART-seq mRNA kit (v4) (Takara, #634772), with the exception of *NvNcol1*-labeled cells, in which only three replicates were sorted due to tissue availability. For planulae, two replicates of 1,500 cells and four replicates of 100 cells were sorted using the same parameters. Samples were maintained at 4C for the duration of the sort, and after sorting spun down briefly, flash frozen and stored at −70C prior to cDNA synthesis.

#### cDNA Synthesis, Library Preparation and Sequencing

1,500-cell pools and 100-cell pools were used for cDNA synthesis according to the manufacturer’s directions with the SMART-seq mRNA kit. cDNA was amplified using either 14 or 18 PCR cycles for the 1,500-cell pools and either 18 or 20 PCR cycles for the 100-cell pools (see **Supplemental Data S3**). Next generation sequencing libraries were prepared from 0.5-1ng of cDNA according to manufacturer’s directions using the NexteraXT kit (Illumina, #FC-131-1096) and IDT for Illumina DNA/RNA UD Indexes Set A, Tagmentation (Illumina, #20027213).

Resulting short fragment libraries were checked for quality and quantity using the 2100 Bioanalyzer (Agilent Technologies) and the Qubit Fluorometer (ThermoFisher). Equal molar libraries were pooled and converted using the Element Biosciences Adept Rapid PCR-Plus protocol for use with the Element Adept Rapid PCR-Plus Kit (Cat. #830-00018). The converted and quantified pools were sequenced on the Element AVITI instrument with a 2×75bp Cloudbreak High Output flow cell (Cat. #860-00004) using instrument software versions current at the time of processing with the following read length: 10 bp Index1, 10 bp Index2, 76 bp Read1, and 76 bp Read2.

#### Data Analysis and Differential Expression Analysis

Raw reads were demultiplexed into FASTQ format allowing up to one mismatch using Illumina bcl-convert (v3.10.5). Reads were aligned to *Nematostella* NV2 genome with STAR aligner (v2.7.11b). Transcripts per million (TPM) values were then generated using RSEM (v1.3.1). Replicates selected for each cell population group have been deposited under GEO GSE309035 (see **Data Availability**) and are listed in **Supplemental Data S3**. Replicates were aligned to the scRNA-seq dataset using Seurat’s CCA integration method. Differential gene expression analysis was performed using edgeR (v3)^69^ with a FC cutoff of 1.5 and FDR cutoff of 0.05. The analysis followed the generalized linear model framework with quasi-likelihood F-tests (“glmQLF”), which provides robust statistical testing for RNA-seq count data for each comparison group. Representative MDS plots are shown in **Supplemental Fig. S10.** A summary of the results for this analysis and annotations based on DIAMOND *blastp* (v2.1.6.160)^70^ searches against UniProt/SwissProt, UniProt/TREMBL (E-value < 0.001), and NCBI nr (downloaded August 2025) (E-value < 0.00001), and *hmmsearch* (HMMER, v3.3.2)^71^ against the PFAM database (downloaded August 2025; E-value < 0.001) are provided in **Supplemental Data S3**. An interactive Shiny app was created to view the edgeR results of each comparison conducted in this study, using annotations from the NV2 genome assembly (see **Data Availability**).

An additional differential expression analysis was conducted using the same replicates with EBSeq (v3.21)^72,73^ through RStudio (v2025.5.0.496 and 2026.01.1+403)^74^ which uses a Bayesian framework to determine differential expression across multiple samples simultaneously. The results with a posterior probability (PP) >0.95 are provided in **Supplemental Data S3**, which also includes biological interpretation of the “Patterns” from the analysis, if feasible. Genes with greatest PP > 0.95 for “Pattern 1” are statistically not differentially expressed between samples and shown in **Supplemental Data S3**.

#### Gene ontology (GO) annotation and enrichment analysis

DEGs across both the adult and planulae datasets were analyzed using Interproscan (v5.77-108.0)^75^ using the -*-goterms* flag to determine gene ontology (GO) terms. GO enrichment was performed via clusterProfiler (v4.16.0)^76^ using *compareCluster* function under the following conditions: fun = enricher, pvalueCutoff = 0.05, pAdjustMethod = “BH”, and qvalueCutoff = 0.20. GO IDs were mapped using the names available from the OBO Foundry, release 2026-03-25^77^. Subsequent dotplots were used for visualization via the dotplot function in enrichplot (v1.28.4).

## Figure Preparation

All images and figures were produced and modified using InkScape (v1.4.1 and v1.4.4)^78^.

## DATA AVAILABILITY

Original data underlying this manuscript can be accessed from the Stowers Original Data Repository at https://www.stowers.org/research/publications/LIBPB-2589. This depository will be accessible upon publication. Prior to publication, supporting data are available from the corresponding author upon request. Raw and processed sequencing data produced in this work have been deposited in the Gene Expression Omnibus (GEO) under accession **GSE309035**. *Nematostella* NV2 gene models used in this work can be accessed at Stowers SIMRbase (https://simrbase.stowers.org/starletseaanemone).

Publicly accessible Shiny applications for the adult scRNA-seq atlas (with cnidocyte-associated subset atlas) and output of edgeR results produced in this work are included below:

- ***Nematostella* Adult Whole Animal scRNA-seq** https://simrcompbio.shinyapps.io/scRNAseq_WholeAdult/
- ***Nematostella* RNA-FACS-seq DGE DataTable** https://simrcompbio.shinyapps.io/Nematostella_RNAFACSseq/

## ACKNOWLEDGMENTS

This work was funded by the Stowers Institute for Medical Research and a National Science Foundation Postdoctoral Research Fellowship in Biology (DBI-2208988) awarded to AMLK. We thank the Stowers Invertebrate team for their dedicated care of our *Nematostella* colonies. We are grateful for the FACS service by Kevin Ferro in the Cytometry Technology Center for construction of the adult scRNA-seq datasets as well as sequencing services by Michael Peterson at Stowers for both the scRNA-seq datasets and RNA-FACS-seq. We thank Andrea Moran for her assistance with injections of *Nematostella* embryos that led to the establishment of both transgenic lines. We thank Mark Miller for the illustrations provided on major *Nematostella* developmental stages in Figure 1. We also thank members of the Gibson lab for their feedback and assistance throughout the project, in particular Steffi Williams and Christof Nolte for experimental assistance and discussions throughout this project and Helen Horkan, Kira Marshall, Christof Nolte, and Victoria Sharp for feedback on earlier versions of this manuscript.

