## Supplemental Materials for "Minicollagen expression dynamics reveal a transcriptional program for cnidogenesis in the sea anemone *Nematostella vectensis*"

### SUPPLEMENTARY MATERIALS

PDF contains descriptions for supplementary data, supplementary figures and legends, and supplementary notes.

**Supplementary Data S1-S3**

**Supplementary Figure S1-S11**

**Supplementary Notes**

**Supplementary References**

**SUPPLEMENTARY DATA**

**Data S1: Relevant sequences for this study. Sheet 1)** Primers used to amplify promoter regions for *NvNcol1>eGFP*$^{cnidae}$, *NvNcol1>eGFP* and *NvNcol5>mScarlet-I* as well as genomic locations of each promoter in NV2 genome assembly. **Sheet 2)** Minicollagen sequences provided to Molecular Instruments for probe set construction. **Sheet 3)** Primers used to produce CISH probes, as relevant to Figures 4 and 6.

**Data S2: Analysis of single-cell RNA-sequencing of whole, male adult tissues and cnidocyte-associated subset. Sheet 1)** Quality metrics for three scRNA-seq datasets from 10X Genomics. **Sheet 2)** Annotation of putative clusters in 'combined, 'whole tissue, adult UMAP projection. **Sheet 3)** Marker table (method = MAST) in 'combined, 'whole tissue, adult clusters where adjusted p-value > 0.05. "NV2_GeneID_annot" refers to annotations from the NV2 SIMR gene models, which corresponds to the Shiny app containing corresponding cell atlas. **Sheet 4)** Annotation of putative clusters in cnidocyte-specific subset UMAP projection. **Sheet 5)** Marker table (method = roc) in cnidocyte-specific subset clusters where AUC > 0.7. "NV2_GeneID_annot" refers to annotations from the NV2 SIMR gene models, which corresponds to the Shiny app containing corresponding cell atlas.

**Data S3: Output of differentially expressed gene analysis from edgeR and EBSeq for minicollagen-labeled cell populations. Sheet 1)** Samples used in DEG analysis. **Sheet 2)** Summary of results from edgeR. **Sheet 3)** EdgeR results and annotations from each minicollagen-specific cell population comparison in this study, where FC > 1.5 and FDR > 0.05. **Sheet 4)** EdgeR results and annotations from *NvNcol3+* cell populations from adults and planulae (4dpf) in this study, where FC > 1.5 and FDR > 0.05. **Sheet 5)** Summary of edgeR results for minicollagens across comparisons. **Sheet 6)** GO-analysis results from *clusterprofiler* for DEGs up-regulated in nematocytes (*NvNcol1+* and *NvNcol6+* cell populations) and spirocytes (*NvNcol5+* and *NvNcol0+* cell populations). **Sheet 7)** EBSeq results and annotations from each minicollagen-specific cell population comparison in this study, where the posterior probability > 0.95. Biologically relevant interpretations of specific patterns are included. **Sheet 8)** EBSeq results and annotations for genes with high posterior probability (> 0.95) of no differentially expression across datasets (Pattern 1), suggesting these are universal cnidocyte candidates.

**SUPPLEMENTARY FIGURES**

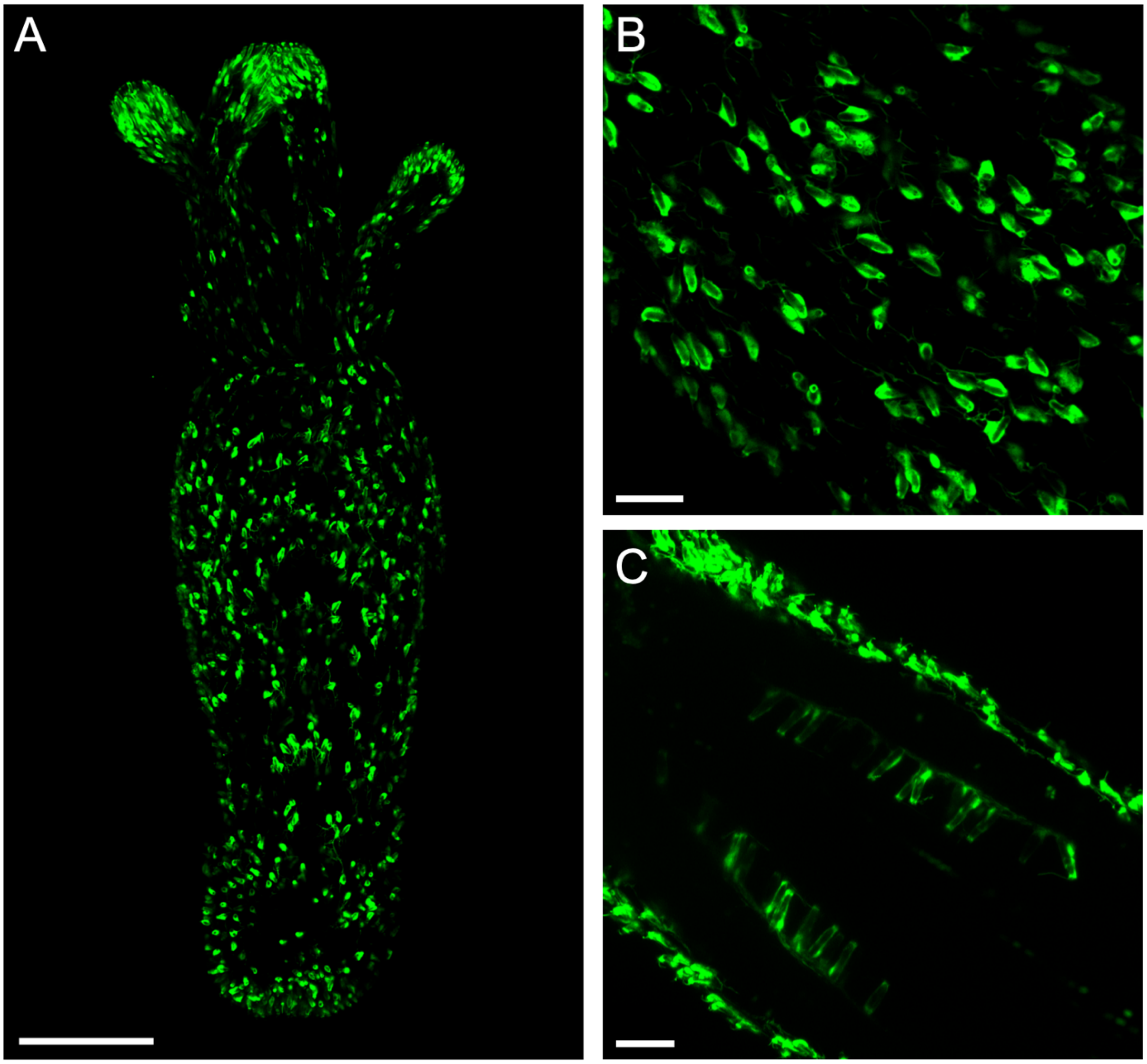


**Figure S1. Live imaging of *NvNcol1>eGFP* transgenic F2 polyp shows eGFP is specific to nematocytes with similar expression patterns as *NvNcol1>eGFP$^{cnidae}$* reporter line.** A) Whole primary polyp. B) Outer ectoderm of body column showcasing developing nematocytes (small isorhizae type). C) Internal nematocytes (mastigophore type) developing along lining of developing mesenteries in central body column of primary polyp with ectodermal nematocytes seen on outer body column. Scale bars: A) 100 µm, B,C) 20 µm.

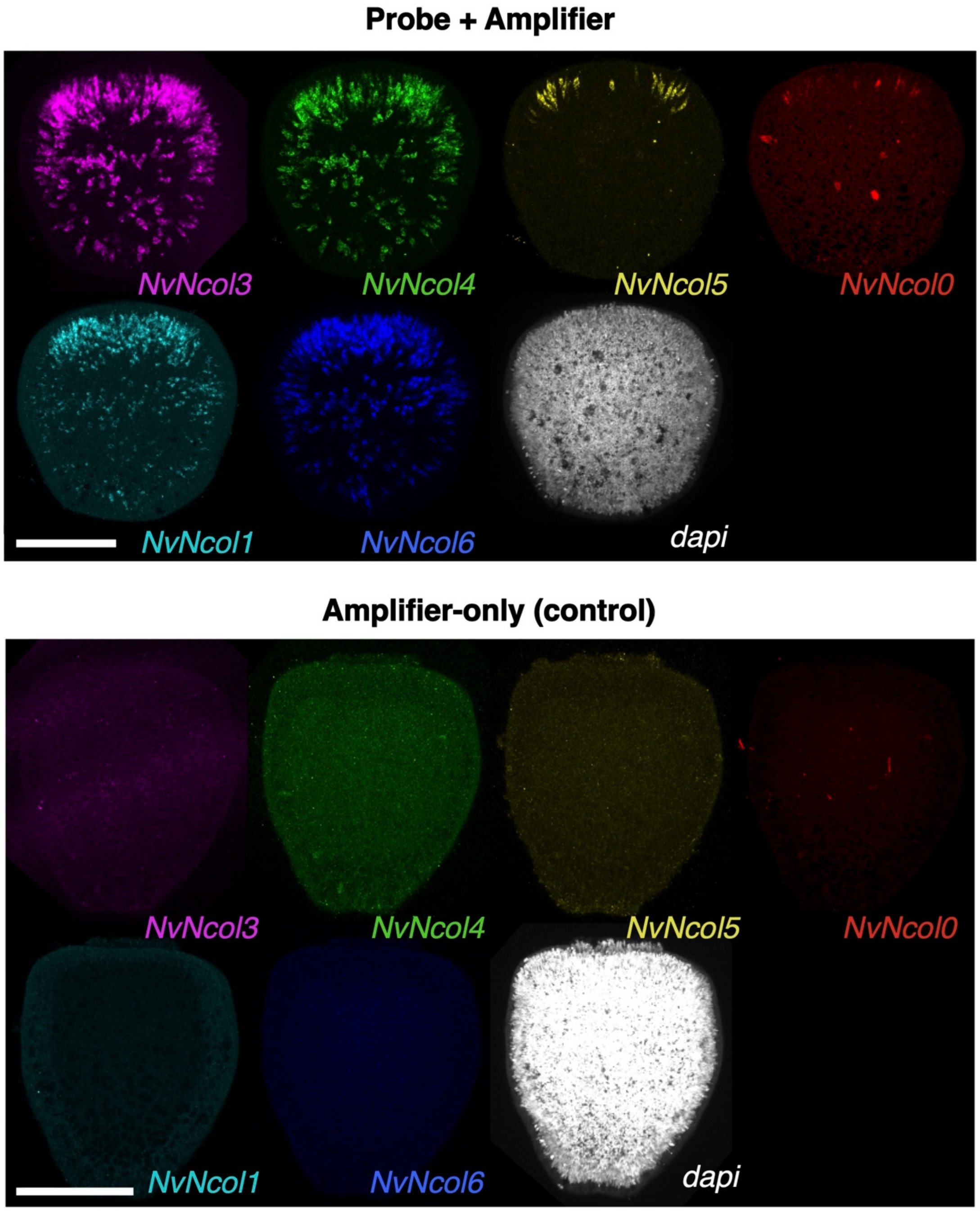


**Figure S2. Representative montage of HCR-FISH experimental and amplifier-only controls for 4dpf planulae.** Note that *NvNcol1* (AlexaFluor 546) and *NvNcol0* (AlexaFluor 555) are split using lifetime unmixing (see Methods). Scale bar: 100 µm.

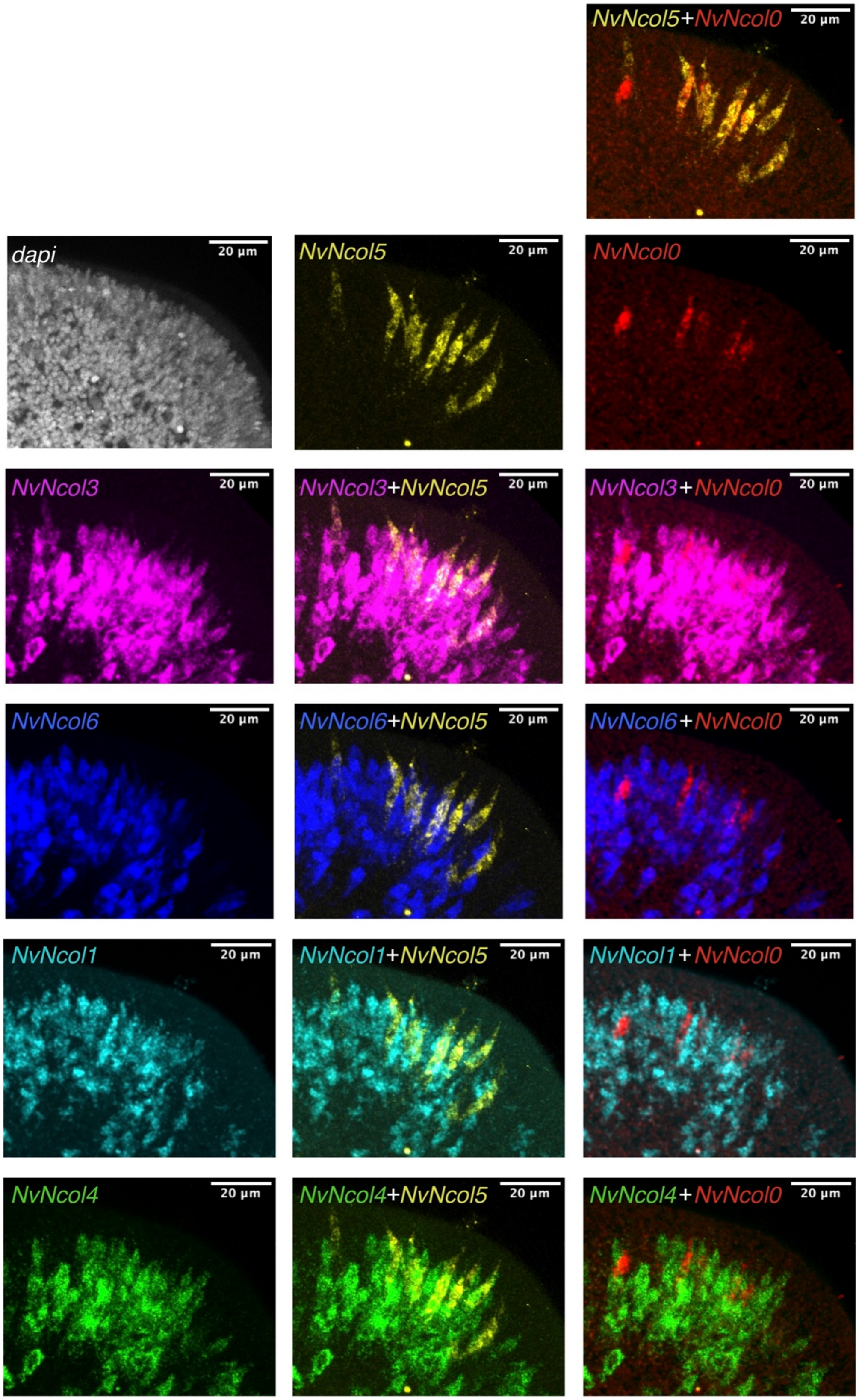


**Figure S3. Overlap of *NvNcol5+* and *NvNcol0+* cells with other minicollagen candidates within budding tentacle of 4dpf planula.** Scale bar: 20 µm.

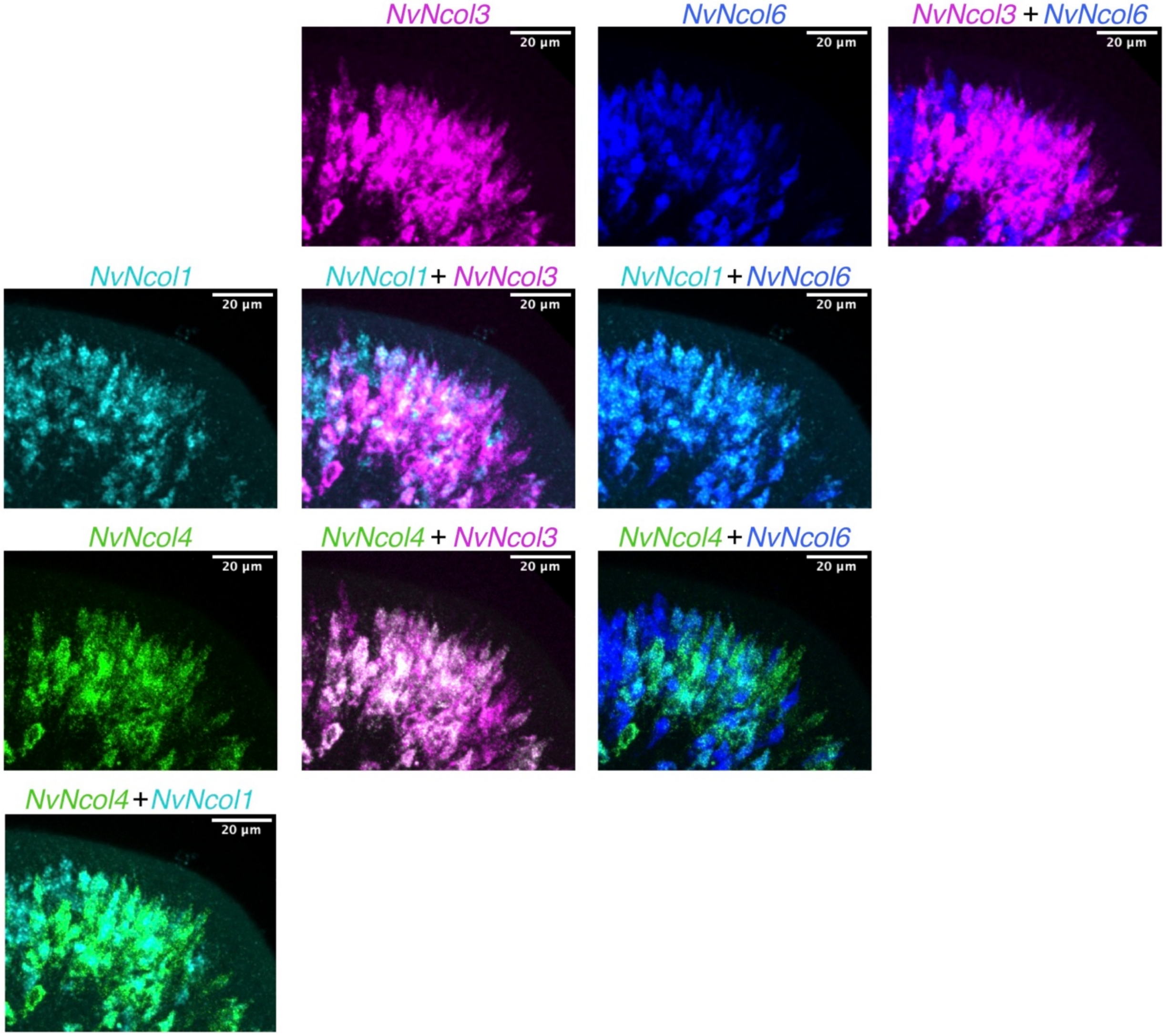


**Figure S4. Overlap of *NvNcol3+* and putative nematocyst-specific minicollagens (*NvNcol1+, NvNcol6+ and NvNcol4+*) within budding tentacle of 4dpf planula.** Scale bar: 20 µm.

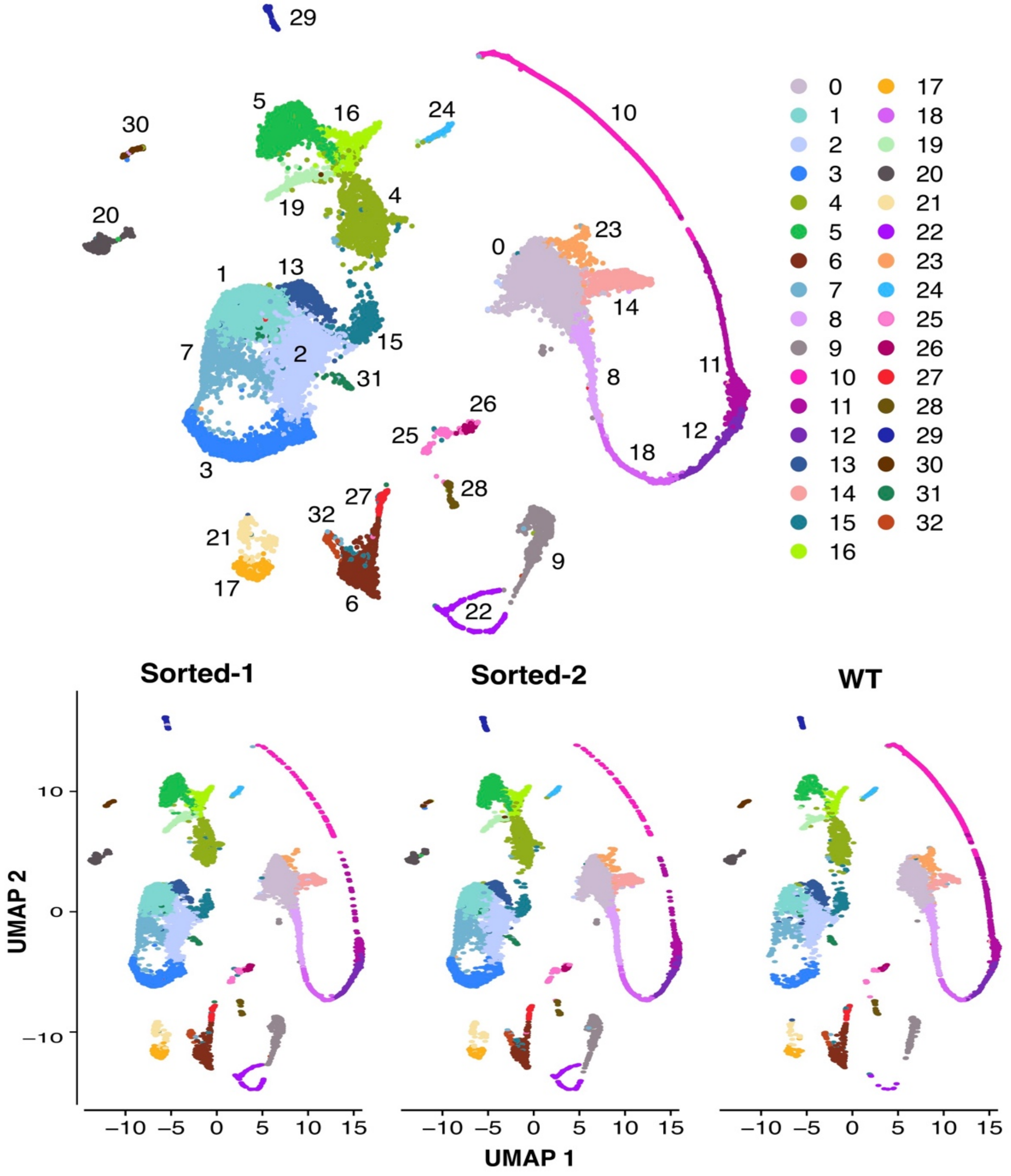


| | # of cells |
|---|---|
| Sorted-1 | 7782 |
| Sorted-2 | 8052 |
| WT | 4892 |

**Figure S5. UMAP for combined, whole-tissue adult male dataset.** Top: Representative clusters in combined UMAP. Bottom: Split from each of the three scRNA-seq replicates, including two replicates from a cell suspension derived from FAC-sorted cells from three individuals of the *Nematogalectin>mScarlet-I* transgenic line ("Sorted-1 and Sorted-2) and one replicate from a separate cell suspension of three wildtype adults ("WT"). Note that all major clusters are represented in each dataset, with enrichment of developing cnidocytes (cluster 22) in the FAC-sorted samples, as expected. Table: Number of cells post quality filtering and used in combined UMAP.

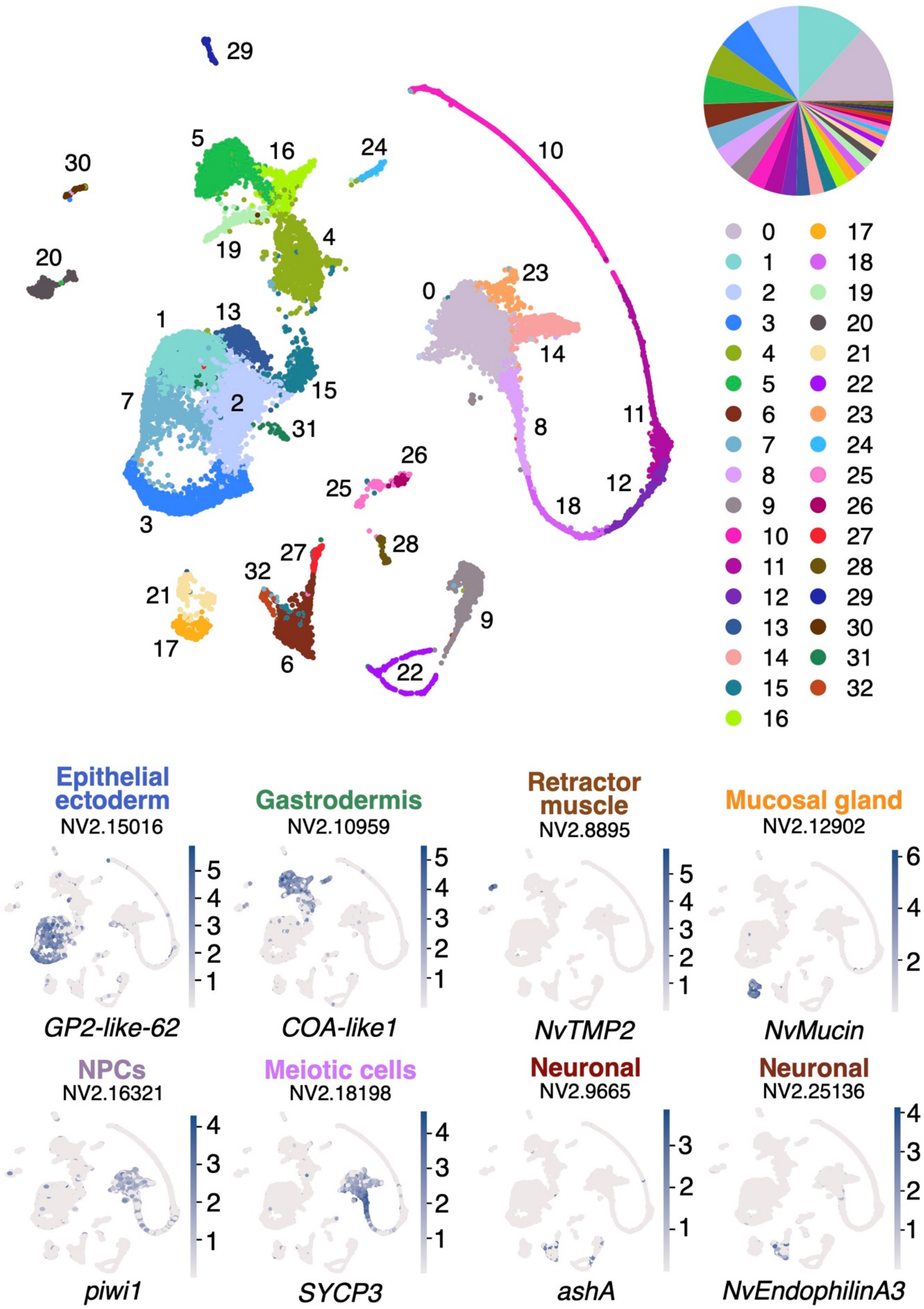

Epithelial ectoderm
NV2.15016
GP2-like-62
Gastrodermis
NV2.10959
COA-like1
Retractor muscle
NV2.8895
NvTMP2
Mucosal gland
NV2.12902
NvMucin
NPCs
NV2.16321
piwi1
Meiotic cells
NV2.18198
SYCP3
Neuronal
NV2.9665
ashA
Neuronal
NV2.25136
NvEndophilinA3

**Figure S6. Representative expression patterns of known cell-type markers in whole-tissue, adult UMAP**. Top: Full UMAP with addition of pie chart indicating proportion of cells in each cluster. Bottom: Representative expression patterns of specific cell-type markers in the whole-tissue dataset (also see **Supplemental Data S2**).

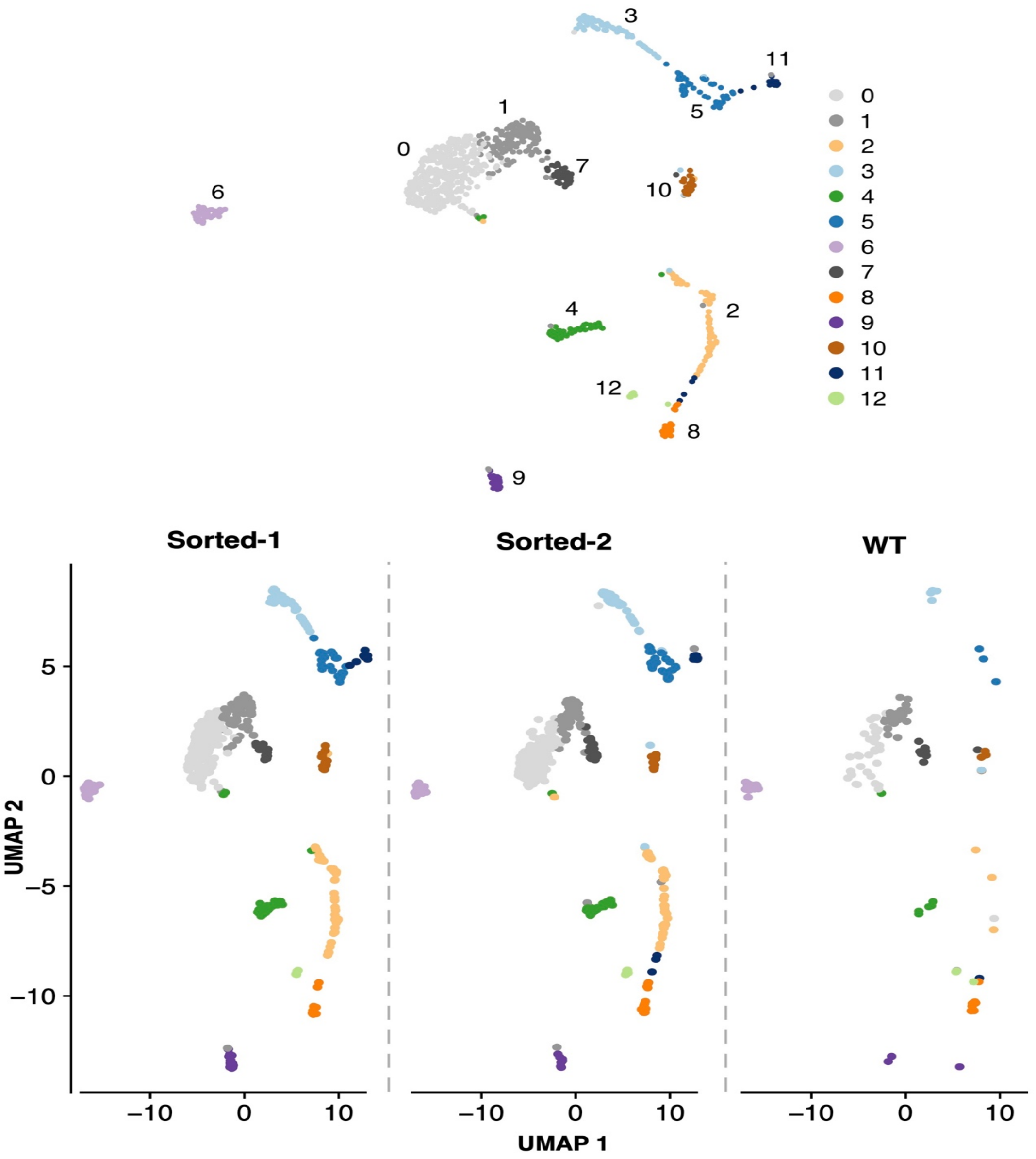


| | # of cells |
|---|---|
| Sorted-1 | 438 |
| Sorted-2 | 422 |
| WT | 124 |

**Figure S7. UMAP for cnidocyte-specific subset of combined adult male dataset**. Clusters 22 and 9 from the combined whole-tissue dataset were isolated and subclustered to produce the above UMAP. Top: Representative clusters in combined cnidocyte-specific UMAP. Bottom: Split from each of the three scRNA-seq datasets, including two samples from FAC-sorted cells from three adults from a *Nematogalectin>mScar* transgenic line ("Sorted-1 and Sorted-2) and one from cell suspension of three WT adults ("WT"). Note significant enrichment of majority of clusters in FAC-sorted datasets. Table: Number of cells post quality filtering and used in combined cnidocyte-specific UMAP.

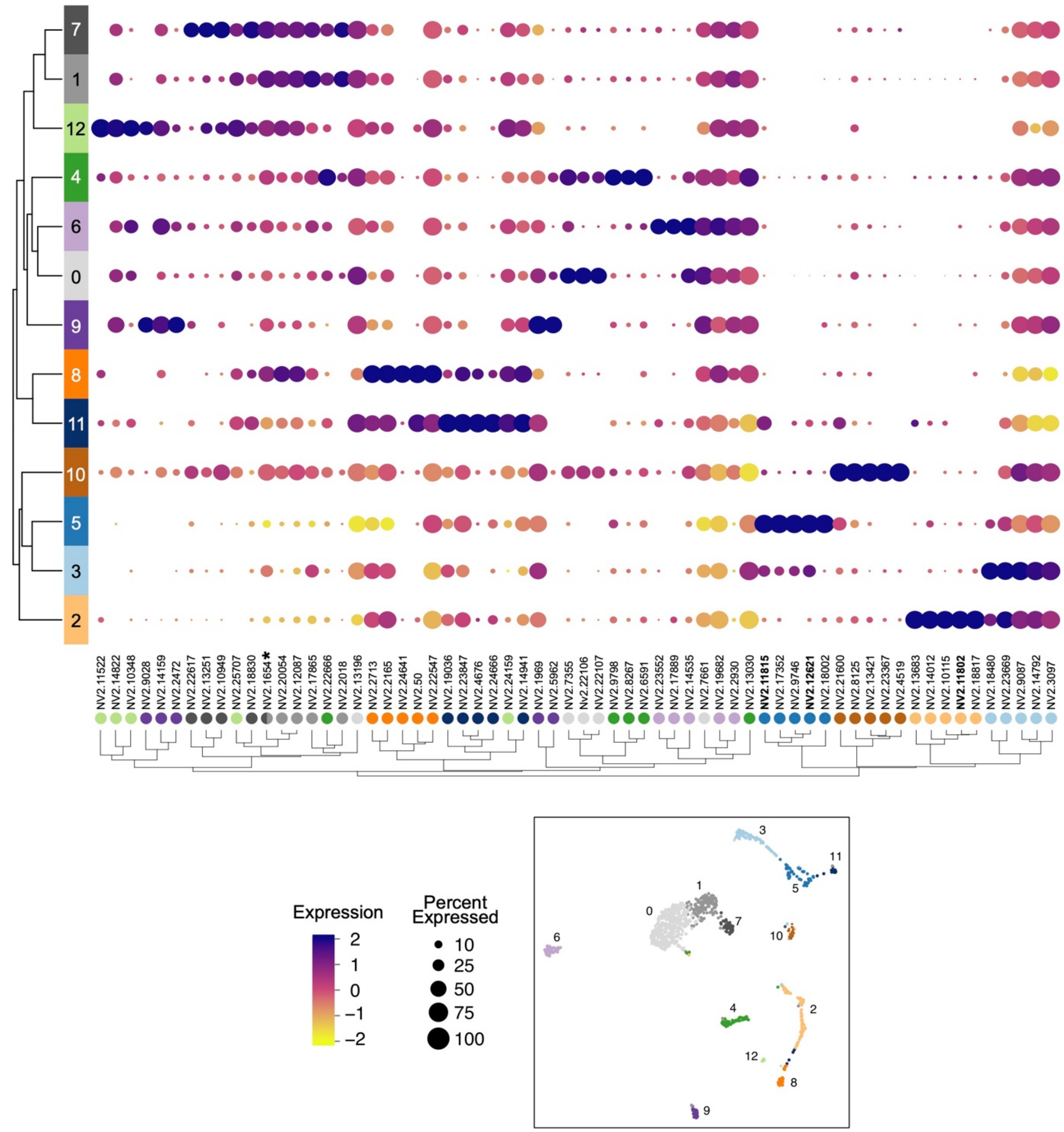


**Figure S8. ClusterDotPlot produced using top five markers for each cluster in the cnidocyte-specific subset.** UMAP of cnidocyte subset shown for reference.

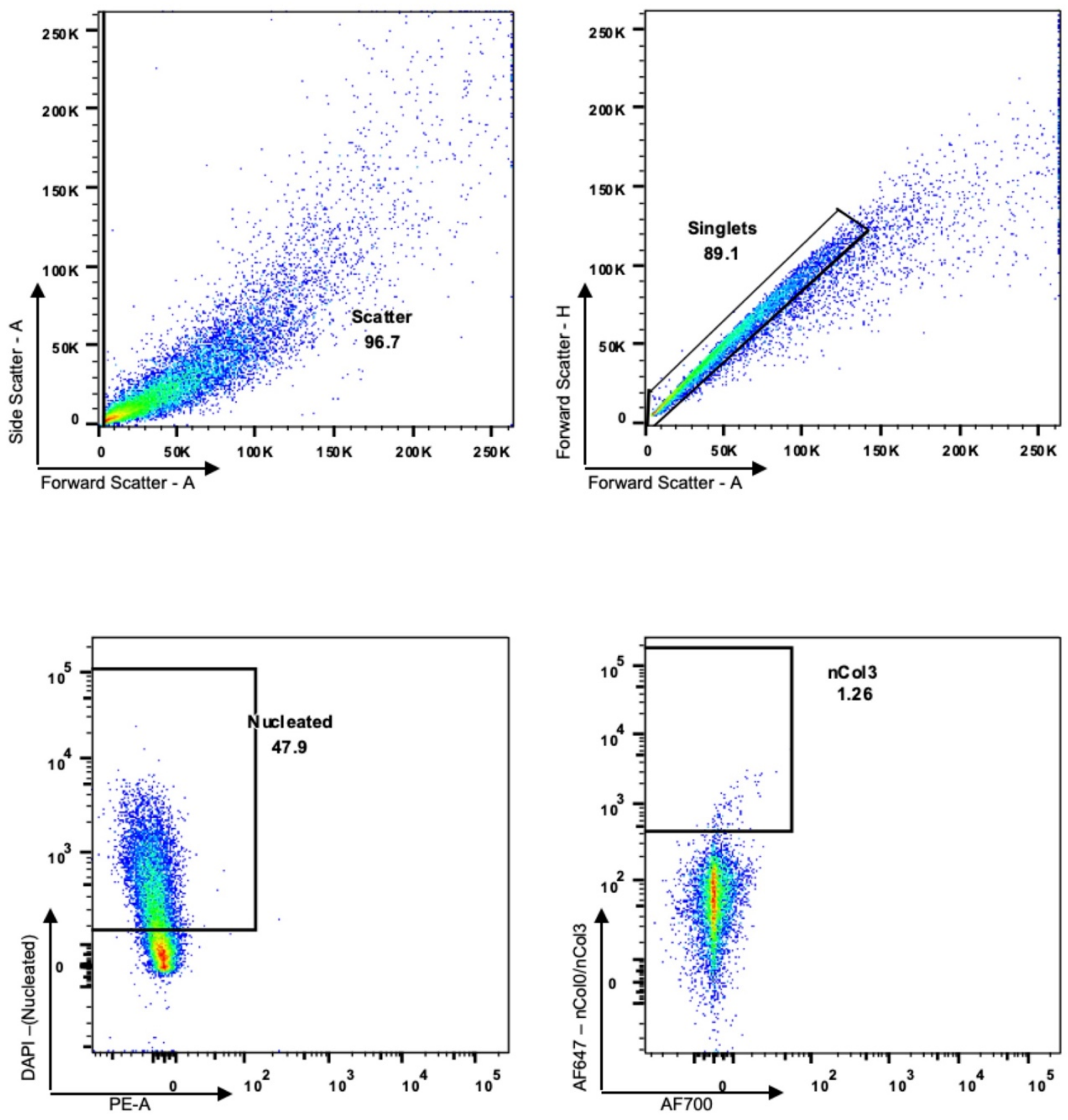


**Figure S9. Representative image of sorting strategy for fixed, HCR-marked samples (here, *NvNcol3+* cells with AlexaFluor 647 amplifier) used for RNA-FACS-seq.**

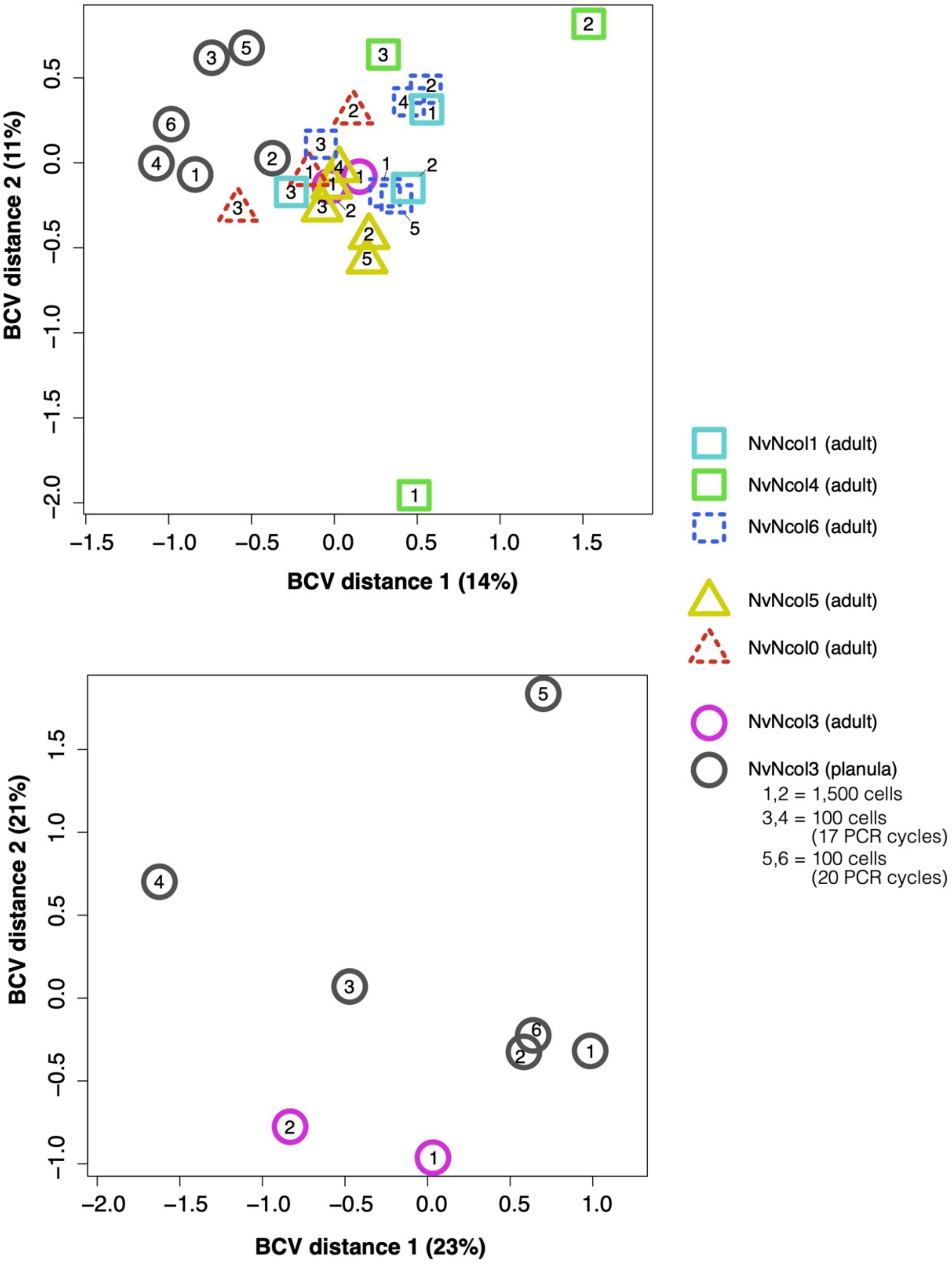


**Figure S10. Representative MDS plot for samples used in differential expression analysis.** Numbers indicate replicate number.

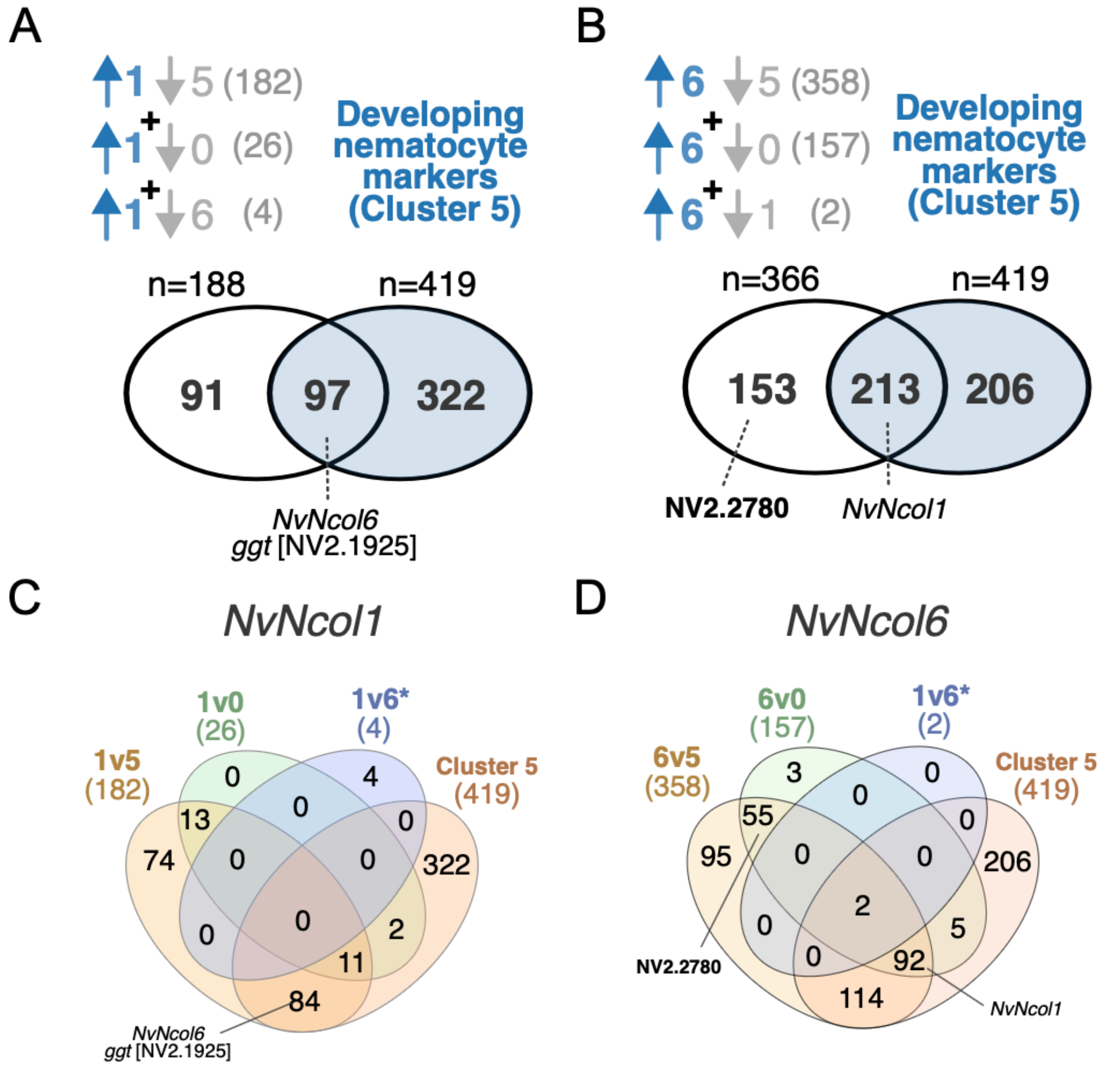


**Figure S11. Venn diagrams showing overlap of differentially expressed genes (DEGs) identified as up-regulated for *NvNcol1+* (A,C) and *NvNcol6+* (B,D) cell populations in comparison to marker genes within cluster 5 of the cnidocyte-specific dataset.** Both *NvNcol1* and *NvNcol6* were identified in the top five markers in cluster 5. In A and B, numbers in parenthesis represent the total number of up-regulated genes in each comparison (e.g. 182 genes were up-regulated in *NvNcol1* cell population compared to *NvNcol5* cell population). Number above white circle in Venn diagram represents non-redundant total of all up-regulated genes from all three comparisons. In A, *Ggt* represents a known nematocyst-specific structural gene. * in C and D indicates pairwise comparisons between *NvNcol1* and *NvNcol6*, which both label nematocytes, thus few genes are differentially expressed between these cell populations.

### SUPPLEMENTARY NOTES

NvNcol0 *and a brief note on names for Nematostella minicollagens*

The original names of the *Nematostella* minicollagens were generated at with the initial genomic assembly[1], which annotated *Nematostella* minicollagens 1-6 based on the order these were located the genome. *NvNcol2* was later found to be a duplicate of *NvNcol1*, and thus was excluded from the study by Zenkert *et al.*[2], which developed the first antibodies for NvNcol1, NvNcol3, and NvNcol4. *NvNcol5* and *NvNcol6* were not included in this study[2]. NvNcol0 is missing a functionally critical domain, i.e. the C-CRD[3-5], thus it may have been excluded as a putative minicollagen in previous annotations of the genome. However, the relatively high and specific expression pattern of this gene was noted in the single cell datasets for adults generated by Sebé-Pedrós et al.[6] and Steger et al.[7], where it was annotated as *Ncol.* Separately, a subsequent study on the matrisome of *Nematostella* named a subsequent minicollagen-like gene NvNcol7[8]. Given the context above, we have annoted this gene as *NvNcol0*.

*Larval* NvNcol3*-labeled cells compared to adult cells in RNA-FACS-seq*

As an additional proof-of-concept for our RNA-FACS-seq methodology, we tested whether adult cnidocytes display different expression dynamics than cnidocytes isolated from earlier developmental stages, namely late-stage budding planulae (4dpf). This is already known to be the case with toxins, such as the larval and maternally-deposited *NvePTx1* (NV2.5238)[9]. We generated six *NvNcol3+* cell populations samples derived from late-stage planulae (4dpf), including two replicates of 1,500 cells and four replicates of 100 cells (**Supplemental Data S3**). The four 100 cell replicates were split into two technical replicates with different PCR cycles, in order to determine the most optimal conditions for reduced cell numbers. We sequenced and compared all six replicates of planulae specific *NvNcol3+* cells with the *NvNcol3+* replicates derived from adults.

We found 784 genes up-regulated in planulae compared to 737 genes up-regulated in adults (MDS plot shown in **Figure S9, Supplemental Data S3**). For example, *NvNcol5* and *NvNcol0* were significantly up-regulated in the adult dataset (**Supplemental Data S3**), which is expected given spirocytes are more abundant in adult tissues (i.e. tentacles). Interestingly, the early life stage-specific toxin, *NvePTx1,* was significantly up-regulated in the planula dataset, despite this toxin being described as a glandular-specific toxin and not produced in cnidocytes. This suggests that there may be some contamination due to the proximity of cnidocytes with *NvePTx1+* gland cells, and this should be taken into consideration when using lower cell input (i.e. 100 cell) samples. However, overall we show that it is feasible to collect as few as 100 cell samples and acquire quality RNA-seq dataset using our optimized RNA-FACS-seq method. Furthermore, this experiment provides multiple genes candidates that differ between larval and adult cnidogenesis for functional experimentation, as the molecular dynamics in adult stages likely display distinct features from early developmental stages.

*Differential expression analysis using EBSeq*

Given our differential expression analysis using edgeR utilize a one-to-one comparative approach, this method is unable to interrogate DEGs that may be up-regulated or down-regulated within multiple populations compared to one or multiple others (e.g. genes up-regulated in both *NvNcol5+* and *NvNcol0+* spirocytes compared to either *NvNcol6+* nematocytes or both *NvNcol6+* and *NvNcol1+* developing nematocytes). Thus, we additionally evaluated this dataset across the six adult cell populations using EBSeq, which compares all experimental samples simultaneously using a Bayesian approach. In total, 79 genes were found to be differentially expressed (posterior probability > 0.95) across the six samples, though with variable expression patterns (**Supplemental Data S3**). An added advantage of this method is that genes with a high probability (PP > 0.95) to not be differentially expressed between samples ("Pattern 1") can also be captured, enabling identification of candidate "universal" cnidocyte-specific markers present across our samples. A total of 3,268 genes matched these criteria as putative cnidocyte-specific genes, which included the known cnidocyte structural gene *NvNematogalectin* (NV2.5200) (**Supplemental Data S3**). These genes may be highly valuable for determined shared gene expression profiles across cnidocyte types, which can be technically challenging using scRNA-seq alone.
