## Supplemental File S1 for "Minicollagen expression dynamics reveal a transcriptional program for cnidogenesis in the sea anemone *Nematostella vectensis*"

**WM-HCR RNA FISH for *Nematostella vectensis***
Modified by Jenny D and Anna K, May 2024

**Reagents*:**

| DAY 0 (Fixation) | DAY 1 (Bleach, digest, hybridization) | DAY 2 (Probe wash, amplification hybridization) | DAY3 (Washing, clearing) |
|---|---|---|---|
| • 7% MgCl2*6H20<br>• **COLD FIX** (0.2% glutaraldehyde/4% PFA/ASW)<br>• 4% PFA/ASW or 4% PFA/PTw<br>• PTw (0.1%)<br>• MeOH | • 3% H2O2 in MeOH<br>• PTw<br>• 4% PFA/PTw<br>• Proteinase K in PTw<br>• Probe Hybridization Buffer<br>• Probe Sets<br>• Salmon sperm (optional) | • Amplification Buffer<br>• Hairpins<br>• Probe Wash Buffer<br>• 5X SSCT (0.1%) | • 5X SSCT<br>• PTw and PBS<br>• Glycerol |

*OF NOTE:*

- *Recipes for freshly prepared solutions in protocol IN BLUE*
- *ASW = artificial sea water (12ppt for* Nematostella*)*
- *Probe buffer, amplifier buffer, and probe solution from Molecular Instruments (v3)*

**DAY 0: FIXATION [~2 hrs]**

Collect animals into clean (RNA-ase free)1.5ml tube (less than 50 ul).
Remove all the liquid
**Cold fix** (see above) for 90s
4% PFA/ASW (COLD) at 4°C for 1 h with shaking
- Alternative: 4% PFA/ASW only (no cold fix) for 1hr either @4C or RT, shaking

Wash 5X with PTw, 5 min @ RT, rocking
Dehydrate tissue using MeOH for, 10-15 min each, rocking, at RT:
- 50% MeOH/PTw, 75% MeOH/PTw, 100% MeOH

Change into new 100% MeOH, store at -20C O/N

**DAY 1: BLEACH, DIGEST, PROBE HYBRIDIZATION [~3h]**

Remove the MeOH from sample
Add 1mL of freshly prepared 3% H2O2 in MeOH to each tube and bleach under bright light for 1hr
- Move tissues to baskets at this time, if using

Re-hydrate tissue with PTw, 5 min, rocking @ RT:
- 50% PTw/MeOH, 75% PTw/MeOH, 100% PTw

Wash 5X with PTw, 5 min, rocking @ RT
Digest tissue by 60 ug/mL Proteinase K for 2 min at RT. No shaking
- Alternative is 20ug/mL for 10 min at RT.
- Prepare ProtK solution during previous washes, keep on ice.

Post-fix samples immediately by 4% PFA/PTw for 30min @ RT. No shaking
Wash 5X with PTw, 5 min, rocking @ RT
Split sample tissue into experimental and controls in 1.5ml tubes

Pellet samples gently and remove PTw.
Add 500$\mu$L of **warmed** (37C) Probe Hybridization Buffer, gently mix with pipette. Avoid bubbles and keep tissue submerged

*Optional for primary polyps: Initial 50% dilution of Probe Hybe rocking at 37C for 10 min. This may prevent collapse of the body column.*

Pre-hybridize for 30 min @37C, rocking
Prepare probe solution in warmed Probe Hybridization Buffer, according to below (adjusting as needed):

| *# Probe Pairs* | *ul of 1mM probes/500ul* |
|---|---|
| *1* | *8* |
| *2-3* | *7* |
| *4-6* | *6* |
| *7-11* | *5* |
| *12-16* | *4* |
| 17-21 | 3 |
| 22-30 | 2 |

Remove 400ul of Hybridization buffer from samples
Add 400 of prepared probe solution, gently mix. Avoid bubbles and keep tissue submerged.

*Optional for primary polyps: Add 50 ug/mL of salmon sperm (10 mg/mL stock solution)*

Incubate O/N @37C, shaking.

### DAY 2: PROBE WASH + HAIRPIN HYBRIDIZATION [~3h]

*Preparation:*
- *Warm Amplification Buffer to RT*
- *Thaw hairpins (protect from light) to RT*
- *Warm Probe Wash to 37C*

Wash 4X with 500ul Probe Wash Buffer, 15 min @37C, rocking

*Optional for primary polyps: Wash 10X with 500ul Probe Wash Buffer, 15 min @37C, rocking*

Wash 3X with SSCT, 5 min @ RT, rocking

*Optional for primary polyps: Wash 5X with SSCT, 5 min @ RT, rocking*

Remove as much SSCT as possible
Add 500 ul of Amplification Buffer, gently mix with pipette. Avoid bubbles and keep tissue submerged
Pre-amplify for 30 min RT, rocking

***For all downstream samples, protect from light***

Prepare appropriate volume of individual hairpins for each experiment in 200ul RNAse-free PCR tubes:
- Use 10ul of each 3mM hairpin stock (**NOTE:** hairpins h1 and h2 must be in individual tubes)
- Snap cool each tubes with following conditions:
  - Heated lid: ON
  - 95C for 90s
  - 23C HOLD

Prepare hairpin solution and add Amplification Buffer to a final volume of 400ul
Remove 400ul Amplification Buffer from each tube

Add 400ul of appropriate hairpin solution, gently mix with pipette. Avoid bubbles and keep tissue submerged

- *Optional: Add DAPI or Hoescht*

Incubate O/N @RT, shaking

**DAY 3: Hairpin Washes + Clearing [~2-3]**

Wash 2X with SSCT, 5 min @ RT, rocking

*For primary polyps: wash 5X with SSCT, 15 min @ RT, rocking*

Wash 2X with SSCT, 30 min @ RT, rocking

*For primary polyps: wash 3X with SSCT, 30 min @ RT, rocking*

Wash 1X with SSCT, 5 min @ RT, rocking

*For primary polyps: wash 5X with SSCT, 5 min @ RT, rocking*

Store in PTw, 80/20 glycerol/PBS, or 90/10 glycerol/PBS

Image as soon as possible, or after O/N clearing in glycerol solution
